# Population dynamics of synthetic Terraformation motifs

**DOI:** 10.1101/095687

**Authors:** Ricard V. Solé, Raúl Montañez, Salvador Duran Nebreda, Daniel Rodriguez-Amor, Blai Vidiella, Josep Sardanyés

## Abstract

Ecosystems are complex systems, currently experiencing several threats associated with global warming, intensive exploitation, and human-driven habitat degradation. Such threats are pushing ecosystems to the brink of collapse. Because of a general presence of multiple stable states, including states involving population extinction, and due to intrinsic nonlinearities associated with feedback loops, collapse can occur in a catastrophic manner. Such catastrophic shifts have been suggested to pervade many of the future transitions affecting ecosystems at many different scales. Many studies have tried to delineate potential warning signals predicting such ongoing shifts but little is known about how such transitions might be effectively prevented. It has been recently suggested that a potential path to prevent or modify the outcome of these transitions would involve designing synthetic organisms and synthetic ecological interactions that could push these endangered systems out of the critical boundaries. Four classes of such ecological engineering designs or *Terraformation motifs* have been defined in a qualitative way. Here we develop the simplest mathematical models associated with these motifs, defining the expected stability conditions and domains where the motifs shall properly work.

## I. INTRODUCTION: TERRAFORMING THE BIOSPHERE

All around the planet ecosystems appear to be experiencing serious threats associated with climate change along with other human-driven impacts (Barnosky 2012, Barnosky and Hadly 2016). Intensive use of land, destruction of regional habitats due to high contamination levels, habitat loss and fragmentation and many other consequences of overpopulation are pushing ecosystems to their limits (Rokström et al. 2009). Some of these systems might be already not far from their tipping point. More importantly, the pace of these responses to external changes is likely to be far from linear, and it has been suggested that it can actually involve discontinuous transitions (Scheffer 2009).

Mounting evidence indicates that even apparently mild, but cumulative changes such as increased grazing, rising temperatures or decreased precipitation can trigger sudden shifts and ecological collapse (Scheffer 2009; Solé 2011). These rapid changes are usually labelled as *catastrophic shifts* (Scheffer et al 2001) typically involving the sudden transition from a given stable ecosystem to a degraded, even fully extinct state. This is the case of semiarid ecosystems. They constitute more than 40% of Earth's land surface and are home of almost 40% of human population (Reynolds et al 2007). Global desertification is a major challenge for our biosphere: current predictions indicate that drylands will expand in the next decades, while some areas can experience rapid collapses. Here minor modifications in external parameters (such as grazing rate) can trigger a rapid decline into a desert state with bare, empty soil unable to sustain vegetation cover (Kéfi et al 2007, Solé 2007). There is now a substantial understanding of past events associated with this type of rapid decline, which is illustrated by the transition from humid, green habitats to bare deserts. About 5500 years ago, the insolation-driven monsoon dynamics experienced a dramatic change, despite the continuous and slow changes associated with insolation and hydrological changes. All available evidence and models indicate that the termination of the green Sahara state was followed by a transition to another stable, alternative state (Scanlon et al 2007). The tipping point found here would then separate two potential attractors (Lenton 2013).

Tipping points are an unavoidable outcome of the intrinsic dynamics of ecosystems and societies (Scheffer 2009; Homer-Dixon 2010; Solé 2011). Due to the nonlinear nature of interactions among species within ecosystems and to the response functions associated with them, the existence of multistability (i. e., the presence of multiple stable states) is the rule, not the exception. For for the same reason, in most cases we can move from one state to another through a “catastrophic” event. Shifts between alternative states are now known to be present in a broad range of situations and have been experimentally demonstrated in micro-, meso- and field scenarios (Scheffer 2007; Dai et al 2012; Lenton 2013). Tipping points have deep consequences for the outcome of the anthropogenic changes of our biosphere. Most policy makers consider the effects of climate change under a risk analysis perspective that somewhat assumes a continuous degradation of the biosphere. Such view is not only wrong, but also highly detrimental for potential solutions or mitigation strategies. We might be running out of time, and different strategies incorporating engineering perspectives might be unavoidable. Geoengineering in particular has emerged as a way of modifying physical parameters (Schneider 2008; Vaughan and Lenton, 2011). This approach includes in particular direct reductions of Earth's albedo through different strategies or the decrease of *CO*_2_ levels through carbon sequestration. None of these strategies is free from staggering costs and long-term efforts and limitations both in their real scale and engineering limitations. What other strategies could be applied in this direction?

It has been recently suggested that an alternative possibility would involve actively changing the biosphere through the use of synthetic biology (Solé 2015). This approach can be used, among other things, as a way to curtail the accumulation of greenhouse gases, remove or degrade plastic debris and other types of waste, act on phosphorous and nitrogen fixation, or slow down ecosystem degradation in arid and semiarid ecosystems (Solé et al 2015; de Lorenzo et al 2016). The key point of this proposal is that engineering living systems allows reaching large scales thanks to the intrinsic growth of living systems, which are capable of self-reproduction. This makes a big difference in relation to standard engineering schemes, where artifacts need to be fully constructed from scratch. Instead, once a designed microorganism is released, appropriate conditions will allow the living machines to make copies of themselves.

This approach, which is an effective way of “Terraforming” the biosphere, needs to consider potential scenarios that guarantee an efficient result as well as a limited evolutionary potential. In this context, target habitats for designed organisms should be chosen as an additional, ecological-level containment strategy. Moreover, limits to the impact of synthetic organisms can be obtained using ecological interactions that are based on either cooperative loops or habitat constraints that are specially well meet by different classes of anthropogenic-modified scenarios. In this paper, we consider a number of possible engineering motifs that can cope with these two constraints. We do not consider explicit case studies (i. e. detailed genetic constructs or designed organisms) but instead the logic design schemes.

As proposed in (Solé 2015) a novel form of addressing the previous issues would be to design synthetic organisms capable of interacting in predefined ways with target species or target substrates in ways that can prevent undesirable responses. The main reason for such approach is that synthetic organisms can be seen as some class of living machine that has been designed as such to perform specific functionalities. The two main differences between these microscopic living machines and man-made counterparts are: (a) they exhibit evolutionary dynamics and thus can change over time as a consequence of mutations and (b) living machines replicate and are thus capable of expanding their populations to scales many orders of magnitude larger than the originally designed populations.

The first difference is something that requires some special attention from the point of view of design. On one hand, evolution will mean in many cases (particularly when dealing with microbes) the loss of the genetic information added to the original organism (Koskiniemi et al 2012), even when the introduced genes involve a fitness gain, as shown for RNA viruses (Willemsen et al 2016). In other words, engineering is continuously needed for a properly executed function. On the other hand, evolving strains that can develop advantageous traits but damage the host ecosystem might create a serious problem. Is there a rational strategy that can minimize the impact of an engineered species? In two previous papers, it was argued that some special classes of engineered ecological motifs might well provide such strategy (Solé 2015, Solé et al 2015). We labelled these basic designs *Terraformation motifs* (TM) since they have to do with our main goal here: to define potential design principles for synthetic organisms capable of addressing ecosystem degradation and climate change.

Our approach to engineering ecological systems requires dealing with multiple scales, as summarized in the diagram of figure 1. Here we consider some levels of complexity, from whole ecological networks and the flows of resources at this level (figure 1a) to the specific nature of interactions among pairs of species (figure 1b) including both wild type strains (*S_w_*) and synthetic strains (*S*) with their hosts (*H*). The upper layers in this scheme already contain engineering components that require considering the cellular networks operating within cells (and how to engineer them, figure 1c) as well as the bottom level description where genetic sequences and available genetic toolkits need to be considered (figure 1d). In this paper, we approach these TMs from the point of view of their underlying population dynamics. We will consider the minimal mathematical models associated with each Terraformation motif and determine the conditions for the survival or extinction of the synthetic organisms. Here too we have a multilevel formal or technical description of all the previous levels (figure 1, right column).

**FIG. 1.**
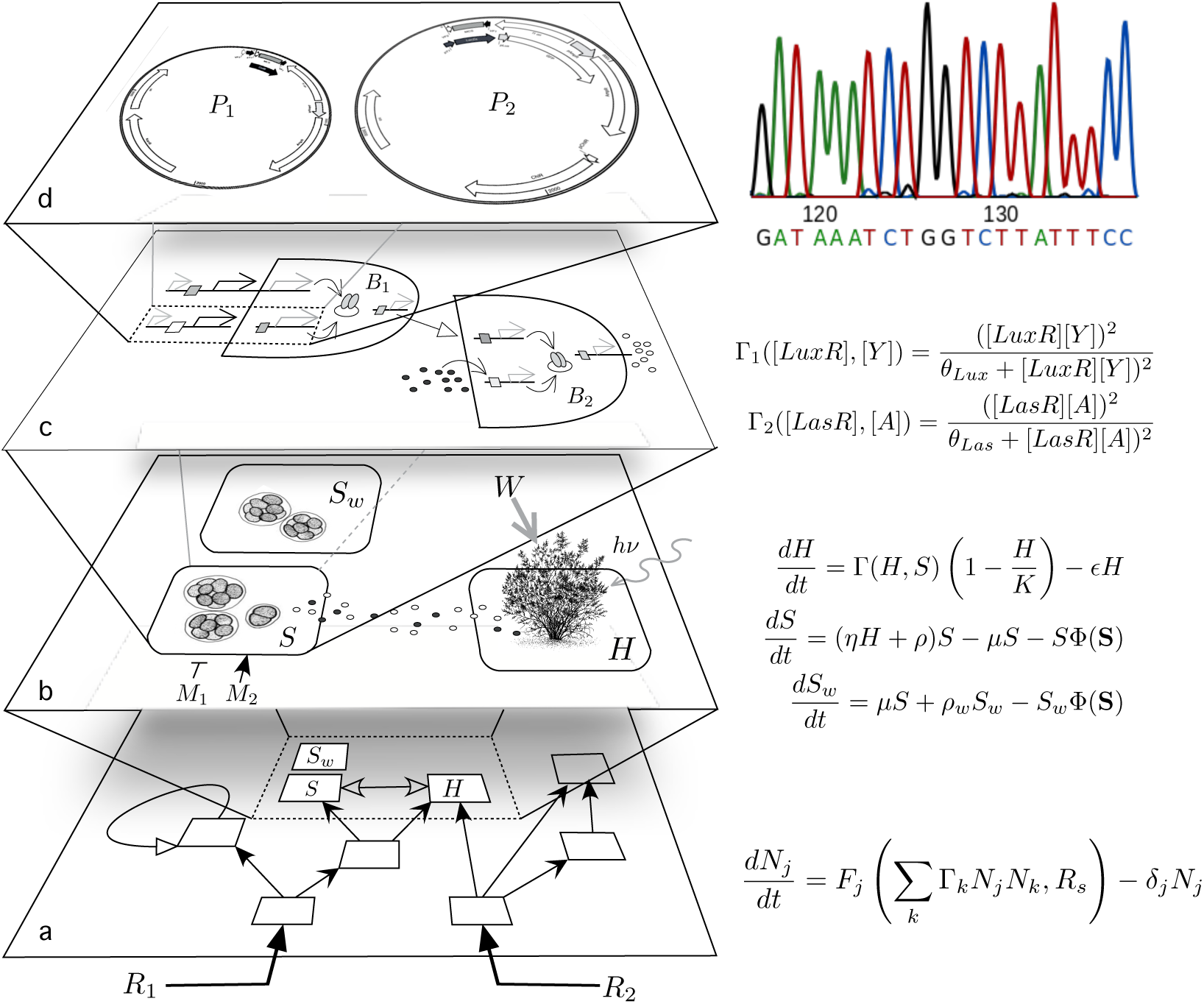
Multiple levels involved in the development of a theory of synthetic Terraformation of ecosystems. Here the different scales are shown at the left and potential mathematical of formal descriptions on the right. Several layers of complexity need to be considered (in principle) including (a) whole population dynamics of ecosystems, with the population levels of each species described in terms of *N_j_* variables. The time evolution of each of these variables would follow a deterministic or stochastic Lotka-Volterra formulation. A smaller scale level (b) considers the dynamics of a few given species, which can include synthetic candidates (here indicated as *S*) derived from a wild-type strain (here indicated as *S_w_*) and a plant host *H*. This small subset can be described, as a first approximation, by means of a small number of coupled equations, defining a three-species subgraph. At the species, cell-level, we also have the mathematical description of molecular interactions typically involving many coupled equations with nonlinear responses among genes and signalling molecules (here we just indicate a typical form of these cooperative interactions terms Γ_1_, Γ_2_). Finally, at the gene sequence level, designed constructs (d) must be engineered in order to operate under predictable circumstances.

There are several reasons why a theoretical approach to these potential synthetic designs needs to be addressed. One has already been mentioned: new and ambitious strategies might need to be developed to solve the problem of catastrophic responses of ecosystems to anthropogenic challenges. Additionally, related problems involving (i) bioremediation applied to highly contaminated areas, (ii) the development of diverse strains of microorganisms to perform useful tasks as symbionts of crops and (iii) ongoing proposals aimed at the control or even elimination of disease vectors (such as mosquitoes) by means of gene drive methods (such as CRISPR/CAS9) imply potential future scenarios where synthetic strains will become incorporated to ecosystems (both natural and novel). Finally, the availability of advanced genetic engineering tools for non-academic groups and the rise of DiY (do-it-yourself) as a parallel avenue for developing synthetic microorganisms calls for a serious effort of understanding the stability and complexity of synthetic ecosystems (Church and Regis, 2012).

The aim of this paper is to make a few initial steps towards a general theory seeking to understand the way these ecosystems might behave and how their populations will achieve different equilibria. Following the scheme outlined in figure 1, two important levels of complexity are presented when studying whole communities (figure 1a) but also when we consider a detailed description of species as cellular networks (figure 1c). As it occurs with standard population dynamics, we need to start with the simple, few-species models indicated in figure 1b, somewhat averaging the details defining each particular partner at the smaller scale and also ignoring the multispecies nature of interaction defining the upper scale. As will be discussed below, such mesoscopic approach makes sense in the contexts discussed here. The models presented below are explicit instances of the four basic classes of Terraformation motifs presented in (Solé et al 2015).

## II. POPULATION DYNAMICS OF SYNTHETIC-WILD TYPE STRAINS UNDER MUTATION

Let us first consider the simplest scenario associated with our Terraformation motifs. It does not define a Terraformation motif in itself, but it does contribute to an important piece of the underlying population dynamics. Here we simply assume the presence of two populations, one being the synthetic organism (hereafter SYN) and the second being the wild type organism(hereafter WT). Here we assume that SYN has been obtained by engineering the WT strain. We also assume that WT can be also obtained back from SYN by mutation or by the loss of the introduced genetic construct (here indicated as *μ*).

### A. SYN-WT mutation model

In this model, the populations of the synthetic strain is indicated by *S* and the population of the WT strain by *S_w_*. Figure 2 displays this system in a schematic way. Figure 2a shows a manipulated organism (microbe) where a given gene has been modified, perhaps adding some additional synthetic constructs either inserted within the genome (such as *S_1_*) or within a plasmid *π* (indicated as *S_2_*). Either case, we simply assume that these constructs can be lost at a rate *μ*, giving place to the WT strain.

**FIG. 2.**
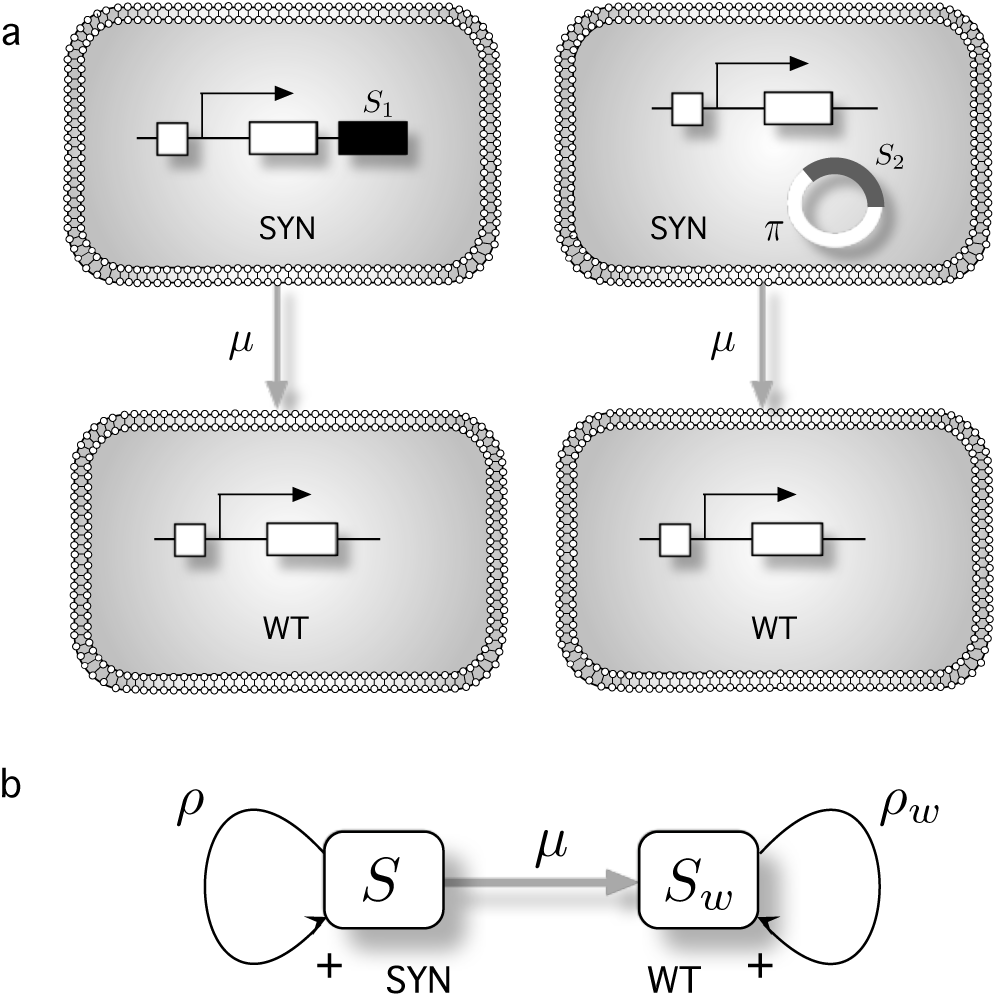
(a) Transitions from a synthetic strain (SYN) towards the original (wild type, WT) species can occur, for example, once the introduced constructs (either within the genome as *S*_1_ or as an external plasmid element *π*, indicated as *S*_2_) are lost. Such a loss of genetic material (the transition SYN → WT) will typically occur at some rate *μ* if the selective advantage provided by the extra genes is too weak. The basic scheme associated with this processes is shown in (b).

The previous processes can be summarised using a transition diagram as shown in figure 2b, where we indicate the population sizes for SYN and WT as *S* and *S_w_* and their replication rates as *ρ* and *ρ_w_*, respectively. If nothing else is included, it is easy to see that the dynamical equations for these two populations are given by:

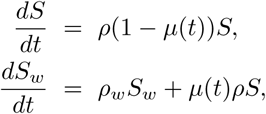

where we have indicated the possibility that reversal to the WT strain can be time-dependent.

Since the first equation is decoupled from *S_w_*, it follows a simple linear growth with *S* and thus gives an exponential form:

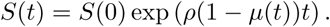

Which now can be introduced into the differential equation for *S_w_* and give:

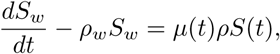

where the RHS of this equation is replaced by the exponential form. The resulting equation can be solved (refs) and give a solution:

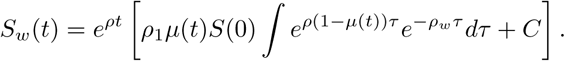

If we assume that *μ* is constant (we leave a more general case for a future study) this general solution gives:

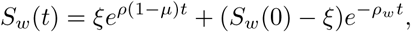

where *ξ* = *μρS*_0_/(*ρ*(1 −*μ*) − *ρ_w_*).

### B. Competition and mutation model

Before we proceed to analyse the three main classes of TMs, let us consider a situation where we simply consider two species, one is the original microbe that inhabits a given environment and its synthetic counterpart, obtained by engineering the first. If nothing else is considered, we can assume that the two strains will compete for available resources. In figure 3 we summarize the structure of their interactions. They compete (as shown by the mutual negative feedback) and also replicate at rates *ρ* (SYN) and *ρ_w_* (WT), respectively.

**FIG. 3.**
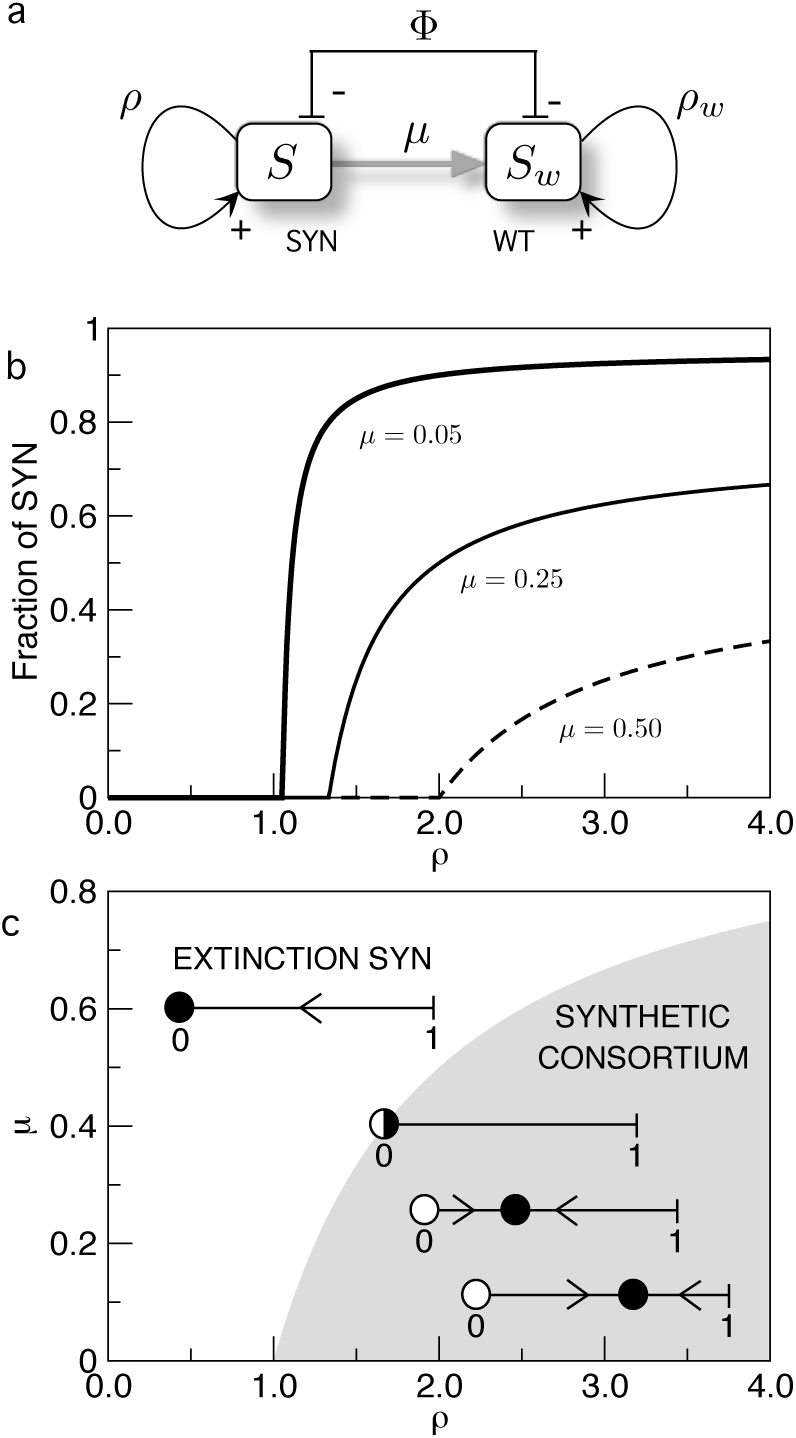
(a) Competition between two strains of organisms where one of them has been genetically modified from the other. Here the engineered synthetic species is indicated as SYN, obtained from an existing one in the same environment, the wild type (here indicated as WT) which can be also obtained back from SYN by mutation (here indicated as *μ*). The populations of each strain are indicated by *S* and *S*_w_, respectively. (b) Bifurcation diagram for three different values of *μ* at decreasing *ρ*, with *ρ_w_* = 1. The fraction of SY N experiences a continuous (transcritical) bifurcation at *ρ* = *ρ_c_*. (c) Phase diagram (*ρ, μ*) displaying the parameter regions with persistence (grey) and extinction (white) of the synthetic consortium. The qualitative dynamics in the one-dimensional phase space given by the line S is displayed as the parameters (from bottom-right to upper-left) approach and cross the bifurcation value (here black circles are stable fixed points, while white circles denote unstable ones).

If only these features are considered, the previous diagram is associated with a dynamical system of competing species described by:

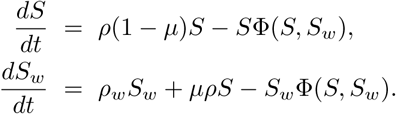

In these equations, we have introduced replication, mutation and competition, as provided (in this order) by the three terms on the right-hand side (RHS) of the equations. Competition is introduced by considering a constant population (see below) and the outflow term Φ(*S*,*S_w_*). In this model, when a given microbe replicates, daughter cells might lose the gene constructs introduced in the engineering process. This occurs at a rate *μ* that gives a measure of the mutation events reverting to the wild type. Such scenario should be expected (and occurs often in experimental conditions) if the fitness advantage of the synthetic organism does not compensate for the metabolic burden associated with the maintenance of additional genetic information.

The previous model (as all of the other models presented below) can be generalized in different ways by considering different functional responses, external inputs, multiple species or stochastic factors. These scenarios will be explored elsewhere. Our interest here is to illustrate the presence of well-defined qualitative dynamical classes of population dynamics. The competition-mutation model considered here can be reduced to a single-equation model if we assume that our species share a given limited set of resources in such a way that their total population *S* + *S_w_* is constant. This *constant population constraint* (CPC), which allows simplifying the previous system, implies:

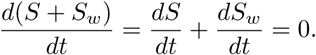

If we introduce this constraint in the equation for the synthetic strain and assume normalization *S* + *S_w_* = 1, it is not difficult to show that the new equation for *S* is:

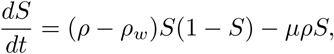

which is formally equivalent to a SIS-like model of epidemic spreading (). As usual, we are interested in the stability conditions associated with the two equilibrium (fixed) points of this system. From *dS*/*dt* = 0, we obtain 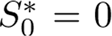 (extinction of the synthetic strain) and the non-trivial fixed point:

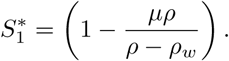

Using linear stability analysis, it is known that a fixed point *S_k_* associated with a one-dimensional dynamical system *dS*/*dt* = *f*(*S*) is stable provided that

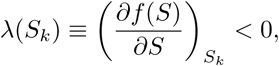

For our model we obtain λ(*S_k_*) = (*ρ*− *ρ_w_*)(1 − *S_k_*) — *μρ*, with *k* = 0, 1. From the previous expression it can be seen that he fixed points exchange their stability when the critical condition *μρ* < *ρ* − *ρ_w_* is fulfilled. In this context, the synthetic organism persists (i.e., 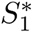 is stable) if:

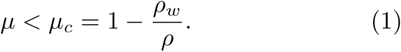

Otherwise, it reverts to WT and gets extinct. This transition is transcritical i.e., the two fixed points exchange stability (the stable one becomes unstable and viceversa) when they collide at the value 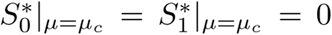 (at the bifurcation point), being 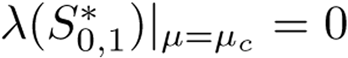. In this sense, when *μ* is increased, the fixed point *S*_1_ moves towards the equilibrium point 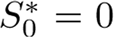, colliding at the bifurcation point *μ* = *μ_c_*. Similarly, from the stability condition Eq. (1) we can derive the critical values of *ρ_c_* and 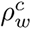, given by *ρ_c_* = *ρ_w_* /(1 - *μ*), and 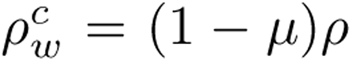. Hence, the transcritical bifurcation will also take place at *ρ* = *ρ_c_* and 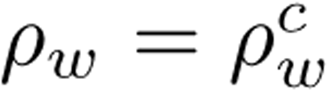. In figure 3b we show the nonlinear behaviour of this system for different values of the rate of reversion. This is obtained by simply plotting 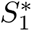 against mutation rate *μ*. Below the threshold, no synthetic organisms are viable, whereas for *μ* > *μ_c_* its population rapidly grows. This means that the competition is sharply resolved once we cross the critical rate *μ_c_*.

In figure 3c we also plot the associated phase diagram by defining the two main phases, using the rate of reversal SYN → WT against the replication rate of the synthetic strain. The persistence of our modified organism will be guaranteed (grey area) provided that it is either enough stable (low *μ*) or enough fast (high *ρ*) in replicating compared to the original strain. Inside figure 3c we display the qualitative behavior of the flows on the line *S* (onedimensional phase space) for each scenario. In the region where the synthetic consortium persists (grey area) the fixed point 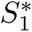 is positive and stable (indicated with a black circle), while 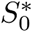 is unstable (white circle). On the other hand, in the extinction scenario (white area), the fixed point 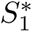 is negative and unstable, while the fixed point 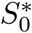 is stable. At the critical boundary between survival and extinction both fixed points collide and become non-hyperbolic (i.e., 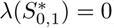).

This type of system is an example of competitive interactions incorporating a mutation term. The main lesson of this model is that a properly designed synthetic organism such that it rarely reverts to the wild type will expand and dominate the system, perhaps removing the wild type. On the other hand, the synthetic microbe must be capable of replicating fast enough to overcome the competition by WT.

Now, we will investigate a slightly different model also describing competition between a synthetic strain and the wild-type. The difference here is that the synthetic strain will contain an engineered genetic construct that can be lost at a rate *μ*. Hence, the mutation term tied to replication will be decoupled from the division of the strain. Now the model is given by:

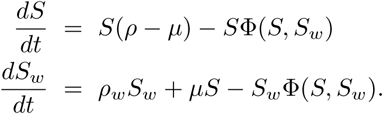

Following the previous procedure we can simplify the two-variable model to a one-dimensional dynamical system describing the dynamics of *S*, given by:

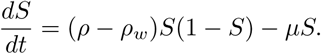

This system behaves like the previous model: there are two equilibrium points that suffer a transcritical bifurcation once value of *μ* is achieved. The fixed points are now 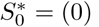, and the non-trivial one,

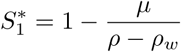

The stability of 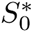 is given by 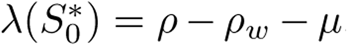, while the stability of 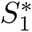 is determined by 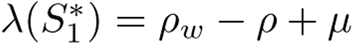.

From the previous values of λ we can compute a bifurcation value of *μ*, given by *μ_c_* = *ρ* − *ρ_w_*. When *μ* < *μ_c_*, *S*_1_ is stable and 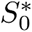 is unstable. At *μ* = *μ_c_* both fixed points collide interchanging their stability since 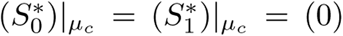 and 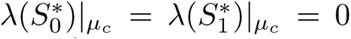 (both equilibria are non-hyperbolic). For *μ* > *μ_c_*, the fixed points 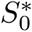 and 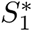 become, respectively, stable and unstable, meaning that the synthetic strain is outcompeted by the wild-type strain.

### III. MUTUALISTIC TERRAFORMATION MOTIFS

An engineered candidate organism to be used for modifying ecological systems should not be capable of decoupling itself from other species in such a way that becomes an expanding invader. One especially appealing scenario is given by engineered mutualistic interactions (figure 4a-b). Mutualism requires a double positive feedback where the synthetic species *S* benefits -and is benefited by- its host *H*. Ideally, design failure should end in the disappearance of the modified species reverting to the wild type. Because mutualism deals with two partners, our synthetic spaces will be constrained by the population of its mutualist partner and such a tight bond is specially convenient, as shown below.

**FIG. 4.**
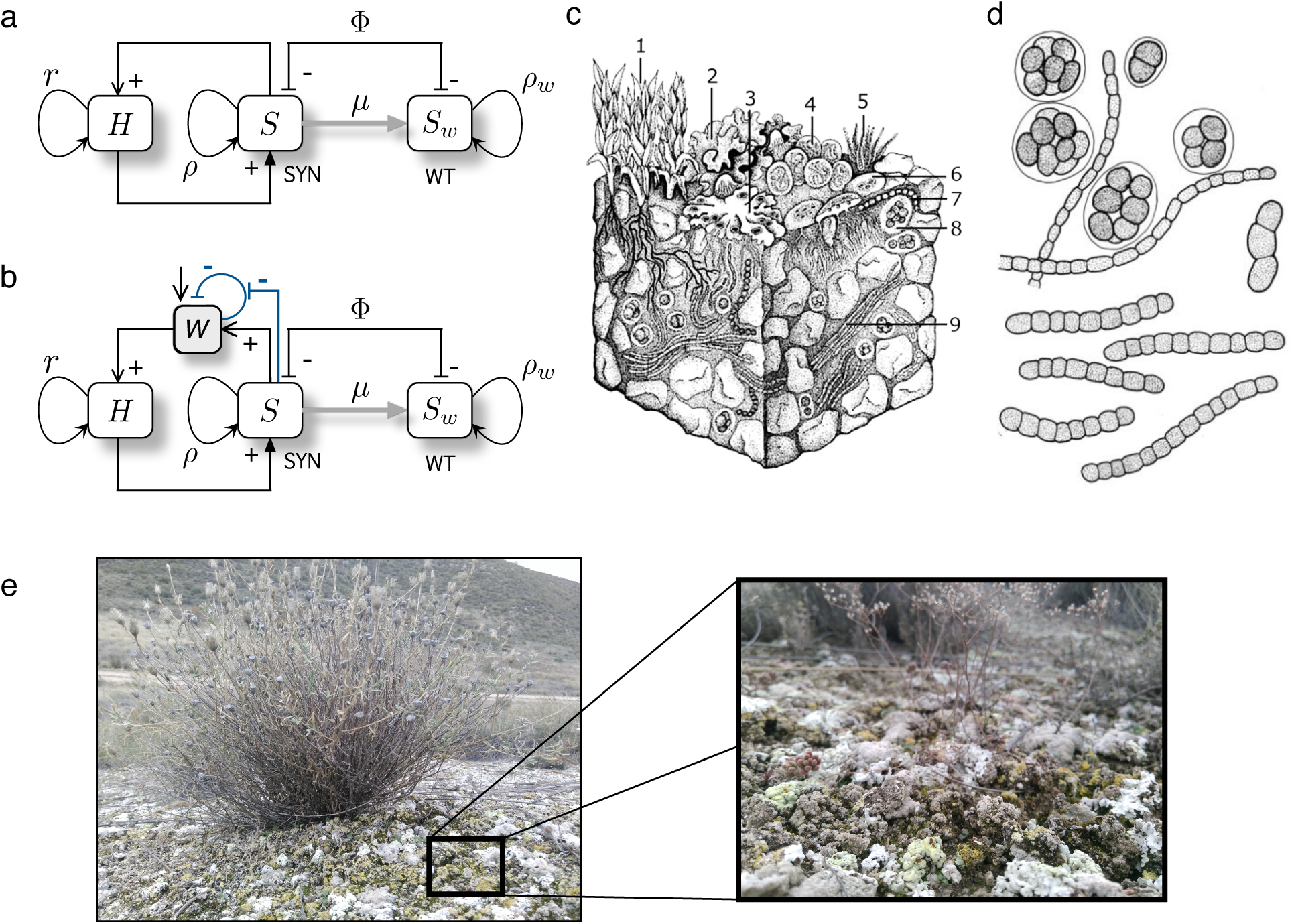
Terraformation motifs involving cooperation among synthetic engineered microorganisms (SYN) and multicellular hosts (H). Considering that the engineered species has been built from a wild-type species (here indicated as WT) living in the environment to be terraformed, the WT can be obtained from the SYN if the engineered construct is lost by mutation (here indicated as *μ*). In (a, b) we display two motifs involving direct (a) and indirect (b) positive interactions among both partners defining a mutual dependency. One potential scenario for this class is provided by dryland ecosystems, where plants would be the host partners and a given local microbial strain the target for the design of a synthetic partner. In these habitats, the soil crust (c) rovides a spatially well-organized community of microbial species that can help engineering cooperative interactions. An engineered microbe capable of improving moisture retention can have a very strong effect on the underlying plant species, expanding their populations. In soil crusts, a whole range of species exist, adapted to water-poor conditions (drawing after Belnap et al 2001). Here we indicate (1) mosses (2,3) lichens, (4,5,7,9) cyanobacteria, (6) fungi, and (8) green algae. An example of these species is shown in (c) where cells belonging to the *Nostoc* genus are represented. (e) Soil crust surrounding an isolated plant in a semir-arid ecosystem from central Spain. The enlarged view displays the detailed structure of the soil crust mainly composed by lichens and mosses.

Several targets can be conjectured. One particularly relevant class is given by the bacteria-root dependencies exhibited by plants and particularly plant crops with their surrounding microbiome. The main case study where this motif applies is provided by semiarid ecosystems, already discussed in the introduction. In these ecosystems, a usually patchy vegetation cover is present, with species adapted to low moisture, extreme temperatures and high UV radiation.

A crucial component of these ecosystems is the biological soil crust (figure 4c-d) defining a complex living skin enclosed within a few centimetres of the topsoil (Weber et al 2016). These are remarkable communities hosting a wide variety of species and largely mediating the energy and matter flows through the soil surface. They are known to help preserve biodiversity and provide a reliable monitorisation system for ecosystem health. In general, the more arid the environment the less diverse is the community, and since plants and the biocrust are strongly related to each other, increased aridity leads to a smaller vegetation cover, less organic carbon reaching the soil, decreased microorganism diversity and reduced plant productivity and a loss of multifunctionality (Maestre et al 2012; Delgado-Baquerizo et al 2016; Maestre et al 2016).

Given that the functional coupling between plant cover and microbial species within the soil crust is already present in these ecosystems, a natural way of approaching a Terraformation scheme is to use the functional links already present. In figure 4 we display the Terraformation motif associated with this engineered design, where, as in the previous example, an extant species (WT) is used as the model organism to build the synthetic strain (SYN). Here we asume that the WT strain does not have a large impact in the plant (our *H* species) but can be engineered in such a way that the synthetic strain *S_w_* is capable of enhancing plant survival.

The basic schemes representing the interactions between the different components of the motif are shown in figure 4a-b. These are of course oversimplified pictures, since we ignore the multispecies composition of the biocrust. This simplification is done with the goal of understanding the behavior displayed by minimal models, in the spirit of fundamental population dynamics (Verhulst 1845, Levins 1969, May 1976). The first case (figure 4a) involves a direct impact through some tight relationship with the host plant, which can be, for example, an engineered symbiosis (Rogers and Oldroyd 2014). The second instead (figure 4b) relies on an indirect cooperation mediated by the influence of the *S_w_* species on moisture. Let us consider and analyse the two scenarios separately.

### IV. DIRECT COOPERATION

The first type of cooperation motif deals with a synthetic strain that enhances the replication rate of the target (host) species. Here the best example would be to start from a free-living species and engineer it in order to built a new strain that becomes an obligate mutualist. Such transition has been shown to be possible and has been created by articially forcing a strong metabolic dependence (Kiers et al 2011; Guam et al 2013; Hon and Murray 2014; Aanen and Bisseling 2014). These studies have shown that the final product can be a physically tight interaction between the two partners.

This case study can be approached by a system of coupled differential equations as follows:

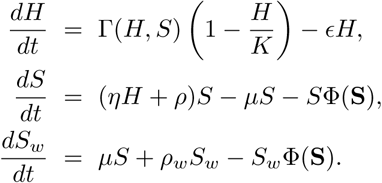

In this model we make the assumption that the two strains (SYN and WT) compete for available space and/or resources while the engineered strain is involved in a cooperative interaction with the host. The state variables of this system are the host (*H*) population, and both the wild-type (*S_w_*) and the synthetic (*S*) strains populations. Here Γ(*H, S*) is a growth function for the host (see below). Parameter *∊* is the density-independent death rate of the host. Constant *η* is the cooperative growth of strain S due to the mutualistic interaction with the host. The other parameters *ρ*, *ρ_w_*, and *μ* have the same meaning then in the previous sections. The Φ(S) function in the of the equations for *S* and *S_w_* stands for the outflow of the system, introducing competition. As we previously did, under the CPC assumption, we can collapse the dynamical equations for the microbial strains into just one, now with:

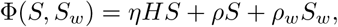

and thus the equation for the synthetic population reads now:

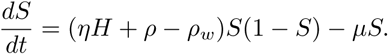

Here we will consider the following form for the growth of the host:

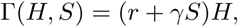

i. e., we assume that the host is capable of growing (at a rate *r*) in the absence of the microbial strains whereas the term *γHS* stands for the cooperative interaction. For *γ* = 0 the host population will grow in a logistic fashion with no direct support from the microbial part. It would simply support it and thus more simple behaviours should be expected. Two potential relevant cases are considered below: a case with strict cooperation where the host can only reproduce when cooperates with microbia, and a case where the host can grow and reproduce with and without cooperation with microbia (i.e., facultative reproduction).

#### A. Strict cooperation

The first scenario considers strict cooperation, which involves that the host can only grow via cooperation (i.e., *r* = 0). For this particular case, the system has four equilibrium points, given by 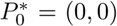,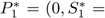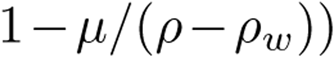, and the pair 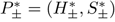 (with *r* = 0, see Secion 1 in the Supplementary Information). The fixed point 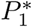 will be outside the positive (biologically meaningful) phase space when *μ* > *ρ* − *ρ_w_*. That is, such equilibrium will only have positive *S* coordinate when *μ* < *ρ* − *ρ_w_*. Under the condition *μ* = *ρ* − *ρ_w_*, the fixed points 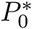 and 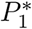 will collide since 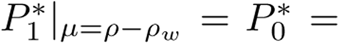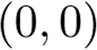. As we will show below, such condition will involve a transcritical bifurcation between equilibria 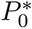 and 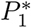.

Let us now study the stability of the fixed points 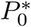 and 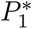 by means of linear stability analysis. Since the expression of the fixed points 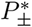 is cumbersome, the analytic derivation of the eigenvalues for these fixed points is rather difficult, and their stability character will be determined numerically by means of phase portraits representation (all of the numerical results presented in this article are obtained by means of the fourth-order Runge-Kutta method with a time step *δt* = 0.1).

The stability of the fixed point 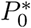 is computed from the characteristic equation 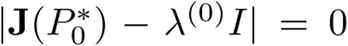, being **J** the Jacobian matrix of the system and *I* the identity matrix. The eigenvalues of this fixed point are given by 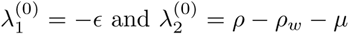. The first eigenvalue is always negative, and thus the stability of this fixed point is given by λ_2_. Hence, the fixed point 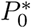 will be stable when *μ* > *ρ* − *ρ_w_*, when *ρ* < *ρ_w_* + *μ*, or when *ρ_w_* > *ρ* − *μ*. Under these conditions this fixed point is stable and thus the host and the synthetic strain will become extinct.

Let us now characterize the stability of the second fixed point, 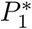. This equilibrium point, if stable, involves the extinction of the host and the survival of the synthetic strain. The eigenvalues computed from 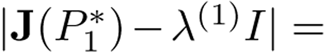0, are given by:

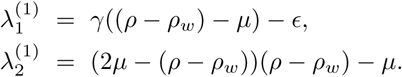

The equilibrium point 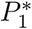 will be stable if

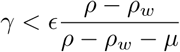

and, additionally, if

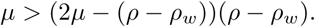

The previous results on the different fixed points and their stability nature are displayed in figure 5. First, we display the dependence of the dynamics in the parameter spaces (*ρ, μ*) (figure 5a) and (*γ, μ*) (figure 5b). Here, for each pair of parameters we solved the system numerically plotting those parameter combinations where H and SYN persist (grey region). Here, there exists a frontier separating the gray and white zones that is given by a saddle-node bifurcation, which creates the pair of fixed points 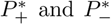, which are a stable node and a saddle. These two equilibria are interior fixed points, and the stable node governs the survival of the host and the synthetic strain. Before the bifurcation, both H and SYN become extinct.

**FIG. 5.**
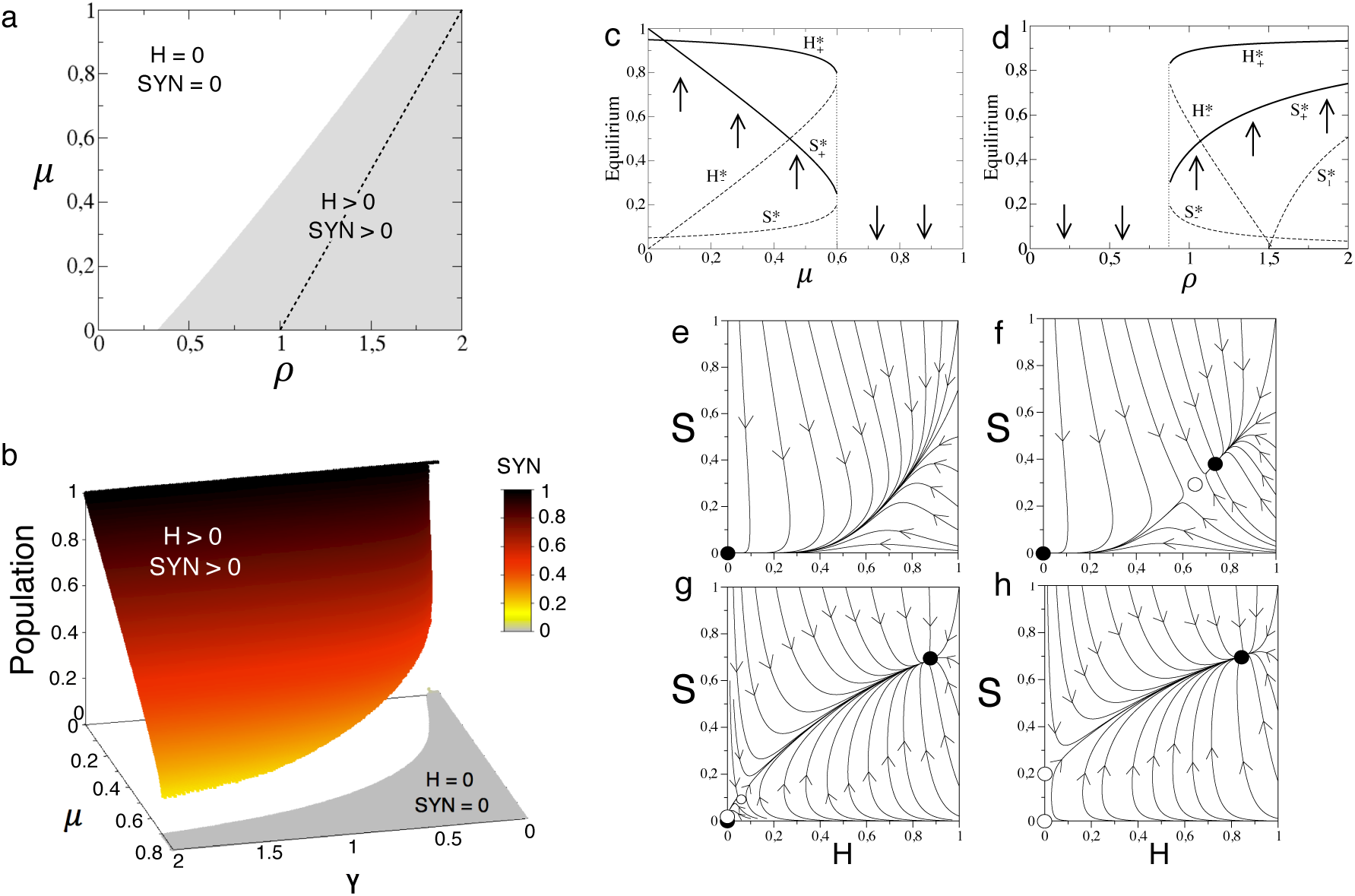
Dynamics of *mutualistic Terraformation motifs* under strict cooperation with *r* = 0, setting *∊* = 0.05 and *η* = *K* = *ρ_w_* = 1. (a) Parameter space (*ρ, μ*) displaying the parameter regions where the host and the synthetic strain survive (gray area) and become extinct (white area), using *γ* = 0.5. The dashed line indicates the transcritical bifurcation. (b) Equilibrium populations for the synthetic strain in the space (*γ, μ*) with *∊* = 0.05, *η* = *ρ* = *ρ_w_* = *K* = 1. In (c) and (d) we display bifurcation diagrams tuning *μ* and *ρ*, respectively, using *ρ* = 1 in (c) and *μ* = 0.35 in (d). In both diagrams we set *γ* = 0.5. Several phase portraits are displayed setting *μ* = 0.5, and: (e) *ρ* = 0.75; (f) *ρ* = 0.91; (g) *ρ* = 1.4; (g) *ρ*; = 1.5. In (f) we use a parameter combination near the creation of the stable node and the saddle after the saddle-node bifurcation. In (g) we use parameter values when the fixed points 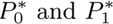 collide. Stable and unstable equilibria are indicated with black and white circles. The arrows indicate the direction of the orbits.

The dashed line in the gray region of figure 5a separates two scenarios where the bifurcation between fixed points 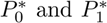 takes place. In the whole gray region the dynamics is bistable, and the system can achieve persistence or extinction of H and SYN, depending on the initial conditions. Figure 5c and 5d shows two bifurcation diagrams by tuning *μ* and *ρ*. The phase portraits of figure 5 display all possible dynamical scenarios with H-SYN extinction (figure 4e), H-SYN coexistence under bistability (figure 5f), the bifurcation between 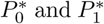 (figure 5g), and the H-SYN persistence without bistability, since after the bifurcation the origin becomes unstable and the node 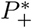 is a global attractor (figure 5h).

#### B. Facultative reproduction and cooperation

The cooperative system considering facultative reproduction of the host (*r* > 0) has 5 fixed points. Two of them are also given by 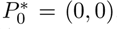, and 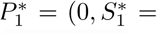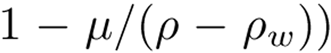, and 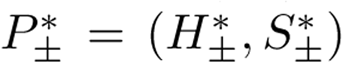 (with *r* > 0, see Supplementary Information), as we found in the previous section. For this system a new fixed point is found, named 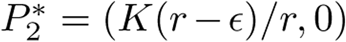. This new fixed point, if stable, will involve the persistence of H and the vanishing of SYN.

The linear stability analysis reveals that the eigenvalues for the fixed point 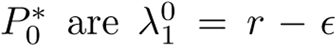 and 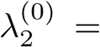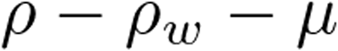. Hence, this equilibrium point will be stable provided *r* < *∊* and *μ* > *ρ* − *ρ_w_*. The stability of the fixed point 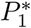 is given by the eigenvalues:

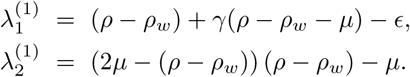

Also, the stability of the fixed point 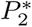 is determined by the sign of the eigenvalues, given by 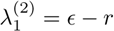 and 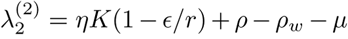. This fixed point will be stable provided *r* > *∊* and

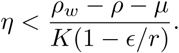

Notice that the stability of this equilibrium also depends on parameters *ρ, ρ_w_*, *μ*, *K*, and *∊*.

The stability of the fixed points 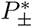 is also characterised numerically, as we did in the previous Section. For the numerical study we will use (if not otherwise specified) a value of *r* = 0.5 > *∊* = 0.05. By doing so we ensure that the fixed point 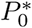 (which, if stable, involves the extinction of both H and SYN) is unstable.

Figure 6 summarizes all the dynamical outcomes of this system. First, we display the equilibrium states of the system in the parameter spaces (*r, μ*) (a); and (*ρ, μ*) (b) computed numerically. The space (*r, μ*) contains three different phases. For those values of *∊* > *r* the outcome of the system is the extinction of H and SYN (the black region in figure 6a), since the fixed point 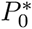 is stable. Two more regions can be identified also in this parameter space. Here, the transition from the scenario *H* = 0 and *SY N* = 0 (black region) to the scenario of *H*> 0 and SYN= 0 is governed by a transcritical bifurcation between fixed points 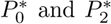 (see phase portraits in figure 6d and 6e). After this bifurcation, the fixed point 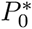 becomes unstable (white circle in the phase portraits of figure 6) and 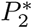 stable (black circle).

**FIG. 6.**
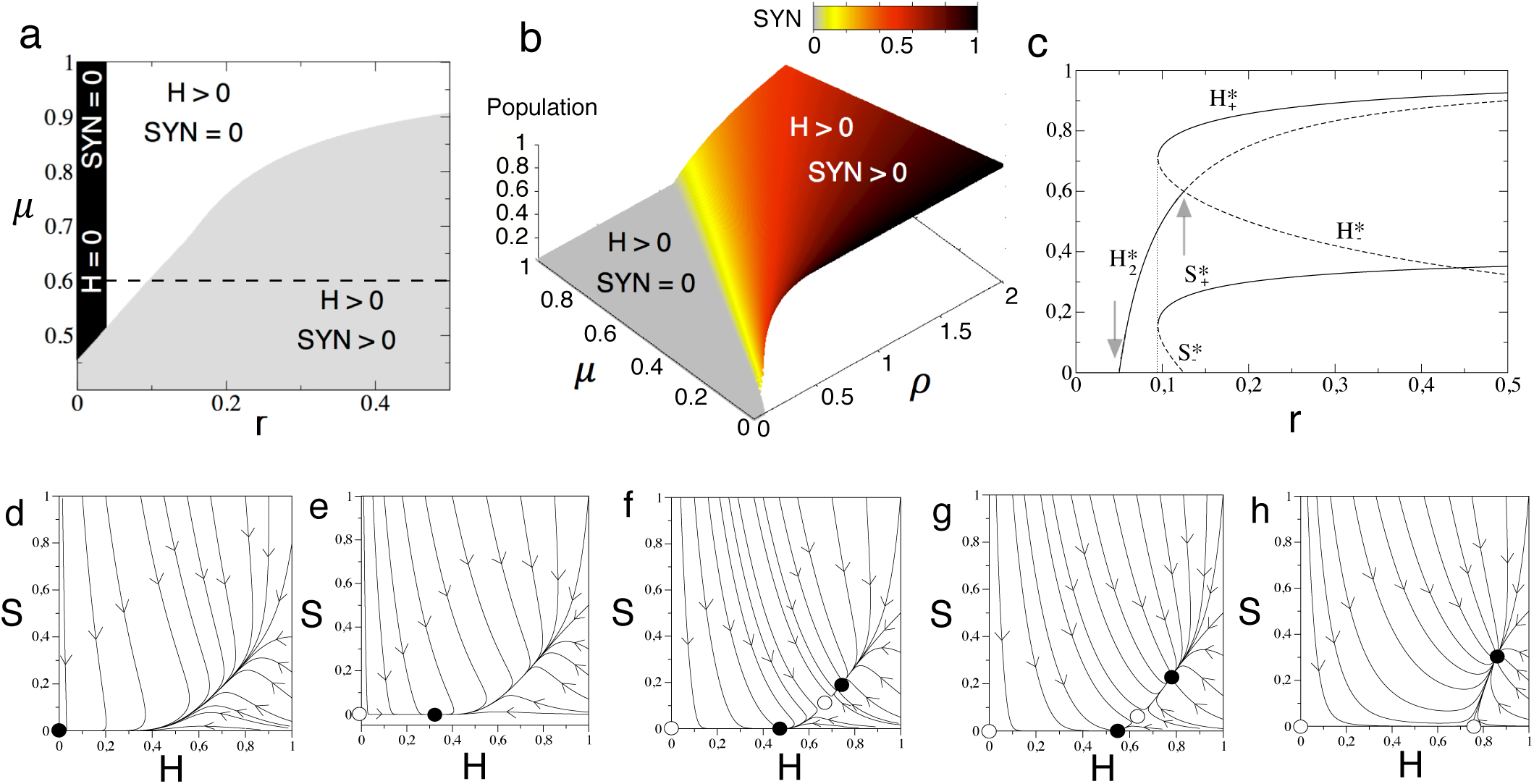
Dynamics and bifurcations for the *mutualistic Terraformation motifs* with facultative reproduction of H, i.e., *r* > 0, using *∊* = 0.05, *γ* = 0.5, and *K* = *ρ_w_* = *η* =1. (a) Phase diagram (*r, μ*) computed numerically setting *ρ* =1. (b) Phase diagram (*ρ, μ*) also obtained numerically using *r* = 0.5. (c) Bifurcation diagram using *r* as control parameter and fixing *μ* = 0.6 and *ρ* =1 (the bifurcation diagram has been obtained tuning the values of *r* following the dashed line in panel (a)). Notice that at increasing *r* the system first suffers a transcritical bifurcation (first gray arrow), then a saddle-node bifurcation (vertical dotted line), and a second transcritical bifurcation (second gray arrow). Phase portraits representation with *μ* = 0.6 and: (d) *r* = 0.045; (e) *r* = 0.075; (f) *r* = 0.0975; (g) *r* = 0.1; (h) *r* = 0.2.

Similarly to the case for strict cooperation studied in the previous section, the boundary of the region where both H and SYN persist defines a saddle-node bifurcation responsible for the creation of the fixed points 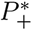 (node) and 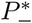 (saddle), and the existence of a bistable scenario. Then, further increase of *r* makes the saddle point to collide with the fixed point 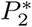 in another transcritical bifurcation. Such a collision involves the interchange of stability between points 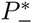and 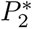. After the collision, the fixed point 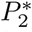 becomes unstable, and the saddle leaves the positive (biologically-meaningful) phase space. Figure 6c displays a bifurcation diagram using *r* as a control parameter. Notice that this bifurcation diagram corresponds to the values of parameter *r* displayed with a dashed line in panel (a) of figure 6 (setting *μ* = 0.6). Here one can follow the series of bifurcations discussed above. Finally, the transition between the white and the grey region in the parameter space (*ρ, μ*) is also given by a saddle-node bifurcation.

### V. INDIRECT COOPERATION

As previously discussed, one of the most obvious candidates to apply the approach taken here is provided by semiarid ecosystems. These and other water-controlled habitats where soil water interacts with a diverse range of soil and community properties, including carbon assimilation, transpiration rates or biomas production (Porporato et al 2002). The biological soil crust (BSC) is composed by a network of mutualists (Bronstein 2016) and provides the ecological context suitable for vascular plants (fig. 4e). It strongly influences key ecosystem processes and its diverse composition offers multiple opportunities for engineering cooperative loops.

Specifically, we aim at describing the impact of an engineered strain capable of improving water retention in the biocrust. This is illustrated by the production of extracellular polysaccharides by cyanobacteria (Mager and Thomas 2011), which have been shown to affect hydrological soil properties, as well as other important features such as soil carbon and maintenance of structural soil integrity. These and other molecules (from vitamins to phytohormones) have been recognised to play a key role in helping plant growth and development (Singh 2014).

The potentially beneficial role of increased extracellular molecules has been exploited in field experiments in desert habitats and sustainable agriculture under adverse ecological and edaphic conditions (Gauri et al 2012). through the direct addition of polysaccharides (Xu 2003), cultivating outdoors combinations of cyanobacteria and plants (Obana et al 2007), and massive inoculation of selected microbial strains (Colica et al 2014). Several positive results have been reported from these studies, suggesting a major role played by ecological interactions connecting molecular, cellular and population responses.

Additionally, the use of

A minimal model that encapsulates the indirect cooperative interactions involving water (state variable *W*) is provided by the following set of equations:

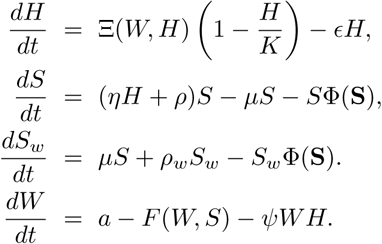

The function Ξ(*W, H*) = *βψWH*, is the growth rate of the host, that depends on the availability of water. Here, the constant *β* is the growth rate of *H* depending on the water, while *ψ* is the fraction of water used by *H* to grow. The population of the host has a logistic growth restriction and a density-independent death rate, parametrized by *∊*. Concerning the dynamical equation for the synthetic strain, constant *η* is the growth rate of the *S* tied to the cooperation with from *H*, *ρ* is the replication rate of the *S*, *ρ_w_* is the replication rate of the WT, and *μ* is the gene construct rate that involves *S* (the engineered organism) to lose function. Now, the dilution flow is Φ(*S*) = (*ηH* + *ρ*) *S* + *ρ_w_ S_w_*.

The last equation includes three terms in the RHS, namely: (i) a constant water input, *a* i.e., rate of precipitation; (ii) water loss, given by the function:

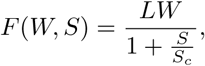

with *L* indicating the maximum evaporation rate and *S_C_* being the rate of inhibition of the evaporation due to the presence of *S*. The function *F*(*W, S*) introduces a specific modulation by means of an inhibition function, namely which includes both the proportionality term *LW* as well as a nonlinear decay associated with the presence of the synthetic population, which is capable of reducing water loss. Finally, (iii) a term of water consumption by vegetation.

As we did for the previous models, we can reduce the system by using the linear relation *S_w_* = 1 − *S*, now having:

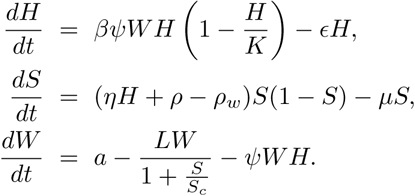

For the sake of simplicity we will use *ρ͂* = *ρ* − *ρ_w_*. This system has five different fixed points,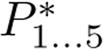, with:

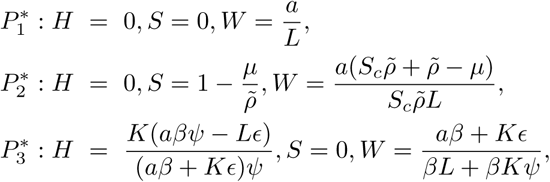

and two more fixed points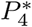 and 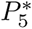 (see Section 2 in the Supplementary information for the values of these fixed points and the Jacobian matrix). The eigenvalues for the fixed point 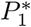 are given by:

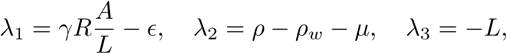

while the eigenvalues for 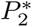 are:

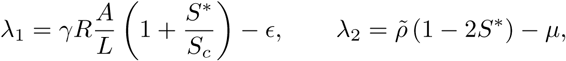

and

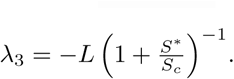

Relevant parameters that could be engineered are the replication efficiency of the *SY N* (*ρ*), the the benefit the SYN obtains from the host (*η*), and the evaporation inhibition due to the action of the synthetic microbia (*S_c_*). Figure 7 displays two-parameter phase diagrams, where the different dynamical scenarios can be visualized. Synthetic organisms will have a higher expression load due to the synthetic construct. This load would make the SYN to grow slower than the wild type (*ρ < ρ_w_*). In order to counterbalance this effect and make the synthetic strain able to survive, the synthetic can take advantage of the vegetation. If there is no symbiosis (*η* = 0) the synthetic organism only survive provided *ρ͂* > *μ* (Fig. 7A). For *η* > 0 the bistable region exists and it becomes bigger as the strength of symbiosis increases. The synthetic survives even for *ρ* = 0 if the symbiosis strength is 1 (Fig. 7B). For higher *η* values the vegetation survives for large evaporation rates (*L*), even with a low replication rate (Fig. 7C).

**FIG. 7.**
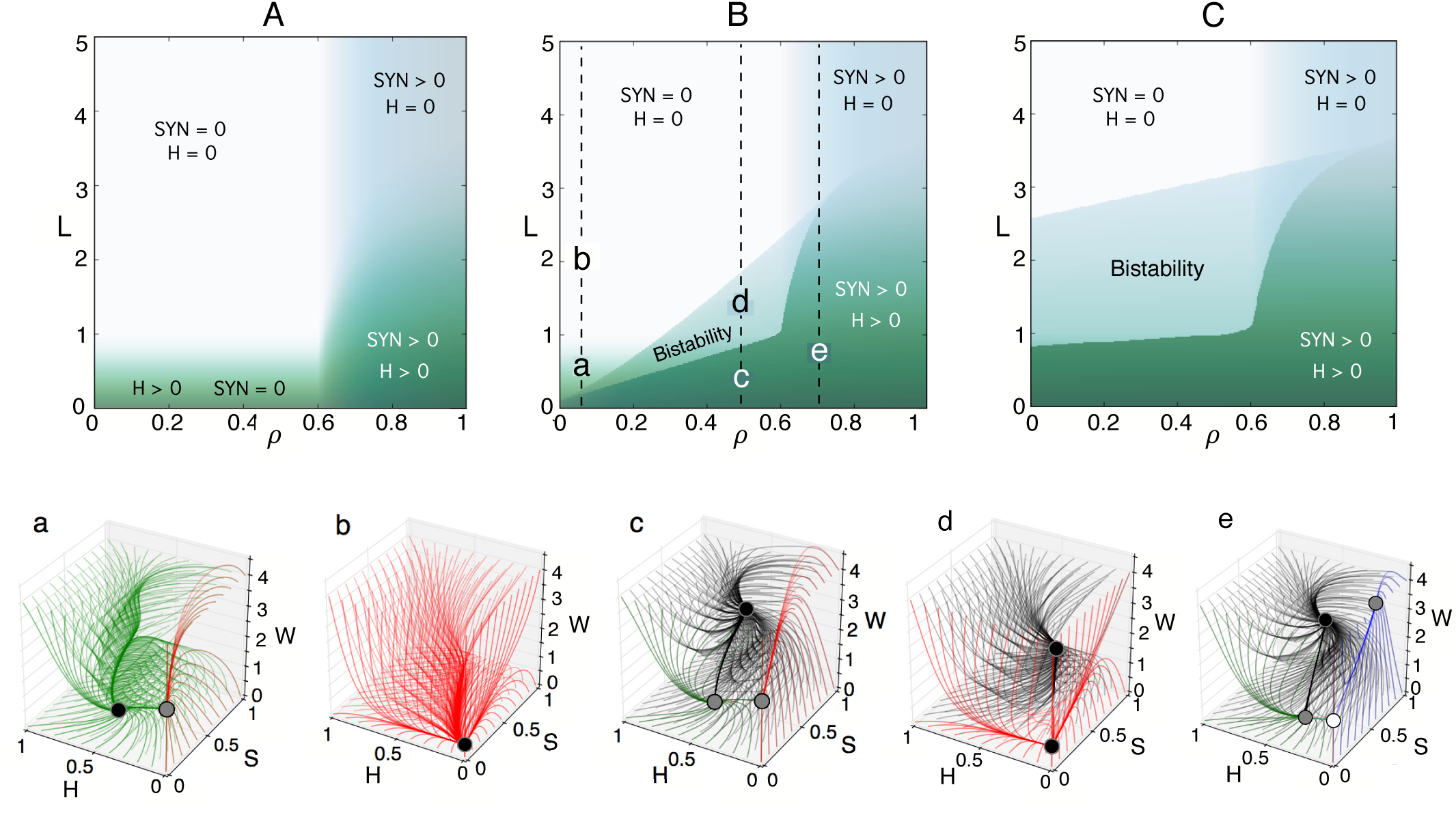
Dynamics for the *indirect cooperation motif* setting: *ψ* = *∊* = 0.5, *β* = *K* = *a* =1, and *μ* = 0.1. (Upper) Phase diagrams displaying the equilibrium for the vegetation (*H*, green gradient) and the synthetic strain (*SYN*, blue gradient) in the parameter space (*ρ, L*) with *S_c_* = 0.3 and *ρ_w_* = 0.5, using three values of *η*: (A) *η* = 0; (B) *η* =1, and (C) *η* = 5. The intensity of the colors for *H* and *SY N* is the sum of the equilibria starting from two different initial conditions (one near to 1 and one near to 0). The vertical dashed lines in panels B and C indicate the parameter ranges used in the bifurcation diagrams displayed in figure S2). (Lower) Phase portraits corresponding to the parameter values indicated with small letters from panel (B) corresponding to the vegetated state (a); desertification (b); coexistence in a vegetated state (c); and engineered bistable ecosystem (d). In case (e) the synthetic strain can survive without the vegetation, and the vegetation can survive if there is not the synthetic strain. The stability of the fixed points is indicated with different colors: stable (black), stable in a plane (gray), and only stable in one direction (white). The color of the trajectories indicates the state reached by the flows: desert (red), vegetated state (green), synthetic and desert (blue), and coexistence between *H* and *SY N* (black).

Nowadays, the semiarid ecosystem is vegetated (figure 7a), but our model reveals that when the temperature rises the system can became a desert (figure 7b). This process and the associated changes in the topology of the phase space can be visualized in *video 1* (Supplementary material). If the temperature is raised in an engineered ecosystem without changing the replication rate, there is a region of bistability, and a sadle-node bifurcation is achieved (see *video 2*). If *ρ͂* is high enough the fixed point 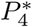 collides with 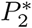 in a bifurcation (this collision can be seen in *video 3*). Once the system is optimally engineered, the vegetation can survive even if *ρ* < *ρ_w_* and the temperature is much higher (figure 7d). With the current temperature, the engineered ecosystem will have the three species (see the fixed point in the interior of the phase space attracting the black trajectories in figure 7d). If *ρ͂* < ρμ the *SY N* will not be able to survive alone (figures 7c and S2a), otherwise the engineered organism will survive even without vegetation (figures 7e and S2f).

The engineering of the system can change the dynamics and the Vegetation-Desert transition from a bifurcation (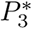 and 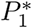) to a sadle-node bifurcation (where both 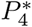 and 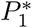 collide). This process can be seen in the bifurcation diagrams of Fig S2. The saddle-node bifurcation involves the emergence of bistability. The bistability leads to hysteresis, meaning that will not be enough to reduce the temperature to recover vegetation (*H*). Nevertheless, if *ρ* is high enough the saddle-node bifurcation take place after a bifurcation of 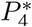 colliding with 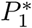. In this scenario, the re-vegetation will take place if the temperature decrease (Fig S2e). The two stable states are the coexistence of the *H*, *S* and *W* and depending on *ρ͂* the desert (figure 7d) or *SYN* and *W* (figure S1g).

Another interesting parameter that could be relevant for a synthetic approximation is *S_c_*. This parameter is related to the effect in the evaporation of water depending on the amount of *SY N*. When *S_c_* is low, the amount of *SY N* population needed to have a positive effect in the environment is very low. This means that with a small fraction of *SY N* the system will be vegetated. This engineered organism could be one that is able to produce a large amount of polysaccharides capable of retaining water. The rate of production will be closely related with the inverse of the threshold. Depending on the difference between the replication efficiency (*ρ͂*) the region of bistability changes. If *ρ͂* < *μ* the bistability region is broader (figure S1A), and if *ρ͂* > *μ* (figure S1B) the coexistence state is more stable. However, the limit where the saddle-node bifurcation take place does not change (figure S1).

### VI. ”FUNCTION AND DIE” DESIGN

Terraformation motifs do not necessarily need to act within natural ecosystems. An alternative scenario, to be considered here and in the next section, is to take advantage of extensive wasteland habitats that have been created by humans and in some sense are already “synthetic”. These synthetic habitats offer opportunities for using bioremediation (including removal of undesired molecules) but can also act as a novel substrates that can host useful synthetic microorganisms.

A synthetic strain could use this substrate as a physical surface allowing it to grow and perhaps disperse. Oceanic plastic debris is an example of this situation (fig 8a). Here a rapidly growing class of new material has been entering oceans and concentrating into large plastic garbage gyres (Jambeck et al, 2015) since the 1950s at a rapid and global scale, leading to the generation of the so called plastisphere (Gregory 2009, Zettler et al 2013). This widespread class of anthropogenic waste is made of non-natural macromolecular structures that were not present in nature and there was thus no biological mechanism expected at that time to degrade it. However, mounting evidence indicates that some species of microorganisms have been adapted to this special class of substrate, effectively degrading it (see Ghosh et al (2013) and references therein) or at least contributing to its fragmentation and decay.

**FIG. 8.**
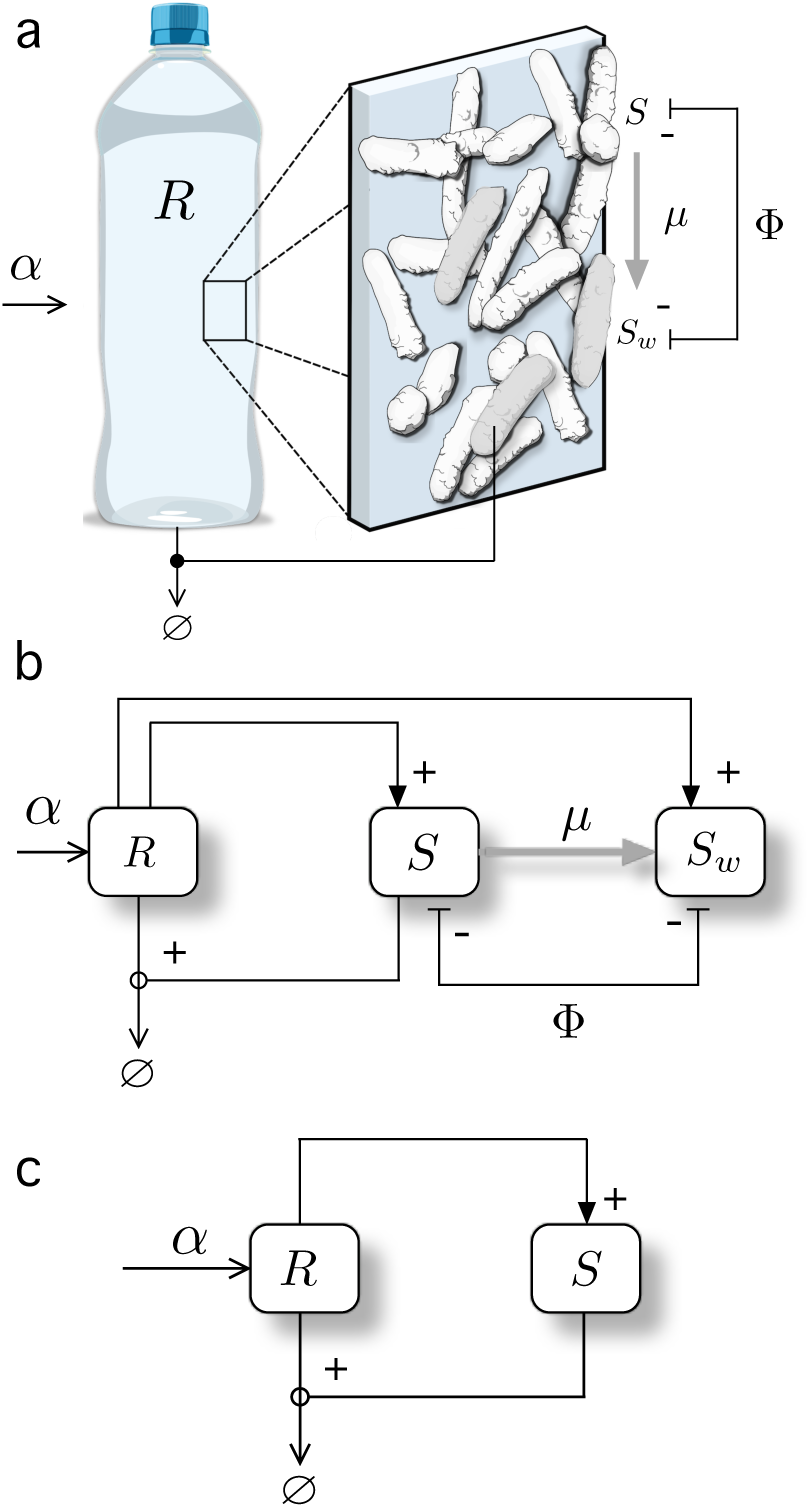
Terraformation motifs for a function-and-die design. This scheme applies to a diverse range of possible targets, such as marine plastic debris (a) where plastic is the resource, entering the system at a rate *α* and spontaneously degraded but also actively degraded by microbial strains *S* and *S_w_* which appear to be supported by the substrate. As in the previous section (and figure), it is assumed that the synthetic strain *S* and its original strain *S_w_* compete for space. The formal motif diagram is shown in (b). In (c) we display a simplified motif where the synthetic strain has not been derived from a wild type variant.

The Terraformation motif analysed here takes its name from an intrinsically relevant property that defines an ecological firewall. The key idea is illustrated by a specific example that has been developed with the goal of repairing (self-healing) concrete cracks, which are a major challenge for the maintenance of infrastructures. The alkaline environment makes difficult for most species to thrive but some species can be used to this purpose. Since repair requires filling a given volume (something that living things can do by reproducing themselves) and do it by means of a suitable material (and bacteria can do that too) synthetic biology appears as a potentially useful approximation here (Li and Herbert 2012).

A microbe can be designed to grow and replenish cracks with calcium carbonate along with a secreted macromolecule that merges into a strong material (Jonkers et al 2010, Rao et al 2013). A major advantage of this problem is that anaerobic bacteria are not going to survive outside the crack and thus selection immediately acts once the task is finished: the function (repair) is done and afterwards the synthetic strain is unable to survive. The right combination of genetic design and ecological constraints create a powerful safeguard.

More generally, we consider here the potential conditions for survival of a synthetic strain living on a given substrate that enters the system and is degraded. The synthetic strain can just degrade the resource or can additionally perform some given functionality. Since removal of plastic debris might actually be part of the goal, it might be unnecessary to use existing species associated with this substrate. Instead, it could be more efficient to simply design or evolve a highly-efficient species capable of attaching to the plastic surface, being also able to outcompete other present species.

The mathematical model associated with the function-and-die motif presented here is given by:

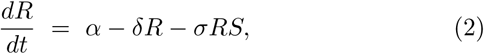

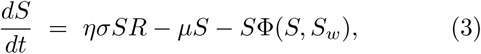

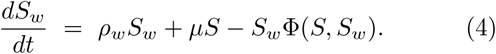

The state variables for this model are the resource (*R*) and both SYN and WT strains. Here *α* is the constant rate of resource (e.g., plastic) income, *δ* is the resource spontaneous degradation rate, and *σ* is the elimination rate of the resource due to the action of the synthetic species. Additionally, *η* is the growth rate of the mutant strain associated with the degradation of the resource (*μ* has already been defined). Finally, *ρ_w_* is the growth rate of the wild-type species.

For this model, the outflow term is given by Φ(*S*, *S_w_*) = *ησSR* + *ρ_w_S_w_*. Assuming again a constant population constraint *S* + *S_w_* = 1, we can see that the equations for the microbial populations collapse into one equation, and the original system can be reduced to the following two differential equations:

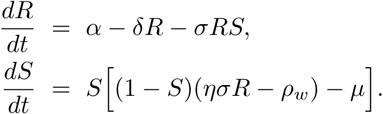

This system has three fixed points, given by

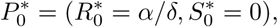

i. e. the only-plastic system) and the pair 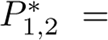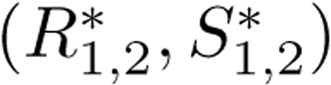 The coordinates of the fixed point 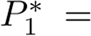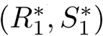 are given by:

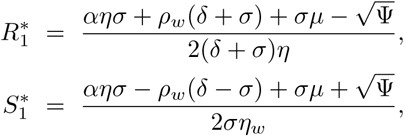

with Ψ = −4*α*(*δ* + *σ*) *ηη_w_* + (*αη* + *δη_w_* + *σ*(*ρ_w_* + *μ*))^2^. The coordinates of the fixed point 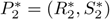 read like the coordinates above but with a change of sign, with:

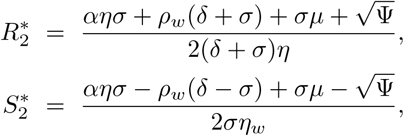

Numerical results obtained for this model suggested that the coordinate 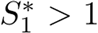 within the range 0 ≤ *η* ≤ 1. For this case, the fixed point 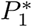 is outside the simplex and it is not biologically meaningful (recall that the CP constraint assumes that 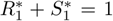, and thus 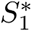 can not be bigger than 1). Under this scenario, the dynamics in the interior of the simplex is governed by the fixed point 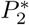. In order to check whether the fixed point 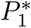 might be inside the simplex, we performed a simple numerical test. We computed the value of the coordinate 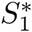for 10^10^ combinations of random parameters (with uniform distribution) within the ranges: *α* ∈ [0, 50], and *δ, σ, η, ρ_w_, μ* ∊ [0, 1]. For all these combinations we obtained values of 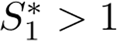.

The stability of these fixed points will be determined by the eigenvalues of the Jacobian matrix,

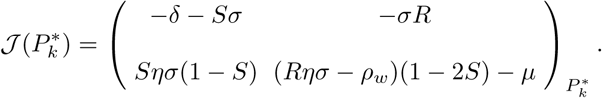

From det 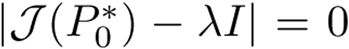, we compute the associated eigenvalues for *P*_O_, given by:

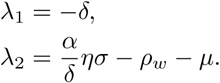

Notice that λ_1_ is always negative and thus the stability of 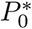 will entirely depend on λ_2_. The change of stability of this point can be computed from λ_2_ = 0. The critical values of the parameters in λ_2_ that involve a change of sign of this eigenvalue are:

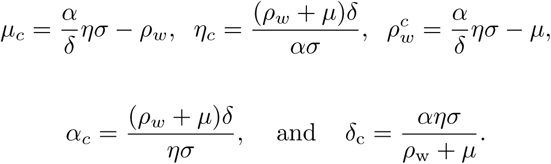

Following the previous critical conditions, the fixed point 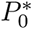 will be unstable (i.e., saddle-point with λ_2_ > 0, meaning that the synthetic strain will survive) when *μ* < *μ_c_, η* > *η_c_, μ_w_* < 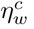, *α* > *α_c_*, or *δ* < *δ_c_*. For example, at *μ* = *μ_c_*, both fixed points 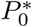 and 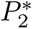 collide since:

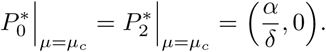

At the bifurcation value these fixed points also interchange stability. Hence, a transcritical bifurcation is found for this motif. The same behaviour is found at *η* = *η_c_*,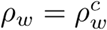, *α* = *α_c_*, and *δ* = *δ_c_*.

Some examples of the bifurcation diagrams associated with this model are shown in figure 9a-b. Here we represent the equilibrium populations *S*\* (computed numerically) of the synthetic strain against the efficiency parameter *η*. A continuous transition given by the transcritical bifurcation takes place for *η* = *η_c_*, when the synthetic strain overcomes the competitive advantage of *S_w_*. Given the definition of *η_c_*, for a fixed input and degradation of the resource and mutation rate, the condition *η* > *η_c_* is achieved once the advantages of the engineered strain overcome the growth rate of the wild type.

**FIG. 9.**
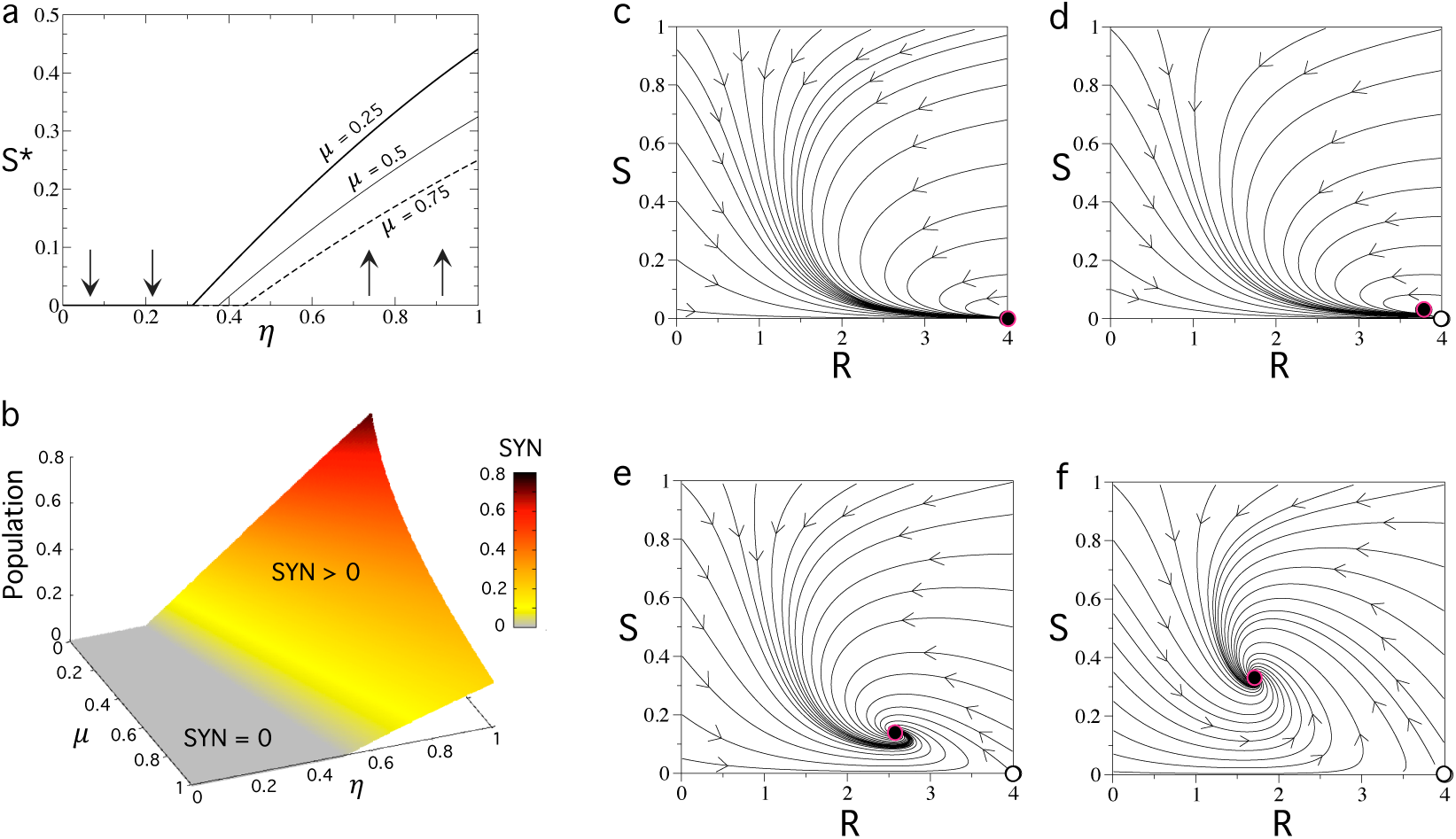
Bifurcations and dynamics of the *function-and-die Terraformation motif* with competing microbes. In (a) we display the stationary population of the synthetic strain (*S*\*) against the parameter *η* that weights the efficiency of the resource-microbe interaction (see text). Here we fixed *α* = *σ* = *ρ_w_* = 1 and *δ* = 0.25, using different values of the reversion parameter *μ*. We specifically use *μ* = 0.25 (thick line), *μ* = 0.5 (thin line), and *μ* = 0.75 (dashed line). The vertical arrows indicate the stability of the fixed point 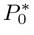. (b) Survival and extinction phases in the parameter space (*μ, η*) computed numerically, using the same parameter values of (a). Several phase portraits are displayed setting *μ* = 0.25 and: (c) *η* = 0.3 < *η_c_* =0.3125, (d) *η* = 0.34 > *η_c_*, (e) *η* = 0.5, and (f) *η* = 0.8. The internal stable fixed point (black circle highlighted in red) is the fixed point 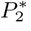. The fixed point 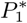 is not biologically meaningful in the range analyzed, since 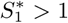 (see figure S3). The unstable equilibrium is indicated with a white circle.

An interesting feature of this diagram is that, even for large values of the reversal parameter *μ* we obtain high population values provided that *η* is large enough. The changes in the phase space at increasing *η* are displayed in figure 9(c-f). In (c) the fixed point 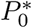 is globally stable (here *η* < *η_c_*. Once *η* > *η_c_* (d-f), the fixed point 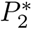 enters into the phase plane having exchanged the stability with 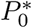 at *η* = *η_c_* via the transcritical bifurcation, thus becoming globally stable. As mentioned, the increase of *η* involves the motion of 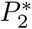 towards higher population values, meaning that the synthetic strain populations dominate over the wild-type ones. The results of the analysis reveal that, provided that the resource is not scarce, we just need a slight advantage of the engineered strain to make it successful and reaching a high population. Moreover, if the resource declines over time, *S_w_* will remain high.

The potential relevance of this scenario is illustrated by the observation that pathogenic strains of *Vibrio sp* might be a major player in the marine plastisphere (particularly plastic microplastic) as revealed by sequencing methods (Kirstein et al 2016). If this is the original (WT) strain, an engineered strain with no toxin genes and improved attachment to plastic substrates could be designed to replace the wild type. Moreover, we should also consider a rather orthogonal scenario where the plastic garbage constitutes an opportunity for synthetic communities to thrive. Since plastic debris is known to be used by a large number of species as a stable substrate, it can be argued that it can be seen as an artificial niche that provides new opportunities for developing complex communities.

It should be noted that plastic degradation by microorganisms is not necessarily good news: degradation (or accelerated fragmentation) of large plastic items leads to a faster transfer towards smaller plastic size, particularly microplastic that can be easily transferred to food webs (Wright et al 2013). Is removal of the human-generated waste a necessary condition for these designed motifs? An alternative Terraformation approach could be using synthetic species that attach to the substrate without actively degrading it. The synthetic microorganism could carry some beneficial function, such as providing useful molecules enhancing the growth or establishment of other species, thus again acting as ecosystem engineers.

Mounting evidence indicates that a rich community of species adapted to these substrates has been developing over the years. Metagenomic analyses indicate an enrichment of genes associated to surface-associated lifestyles (Bryant et al 2016). Within a surrounding environment that is oligotrophic and species-poor, the plastic garbage defines a novel niche that has been fairly well colonised by a wide variety of species attached to the plastic substrate. In many cases, the resulting microbial community provides the scaffold for other species to thrive. Some early proposals on using synthetic biology to address the problem of plastic garbage included a project aimed to facilitate the stable adhesion of plastic pieces with the goal of creating plastic islands. In such a context, we could consider a Terraformation motif where the colonisation by a given species performing other functionalities could be designed, perhaps taking advantage of the niche as an opportunity to build a synthetic ecosystem.

### VII. SEWAGE AND LANDFILL MOTIFS

Our last example in this paper is connected to a major class of waste generated by farming as well as urban and specially mega-urban areas, associated with domestic, municipal and industrial sources. Urban centres incorporate massive infrastructures associated with the treatment of waste as an end part of the city metabolism (Newman 1999). Sewage systems offer a specially interesting opportunity to apply our approach. They contain large amounts of organic matter, along with a wide repertoire of molecules of different origins, from drugs to toxic chemicals. Because of the potential damage caused by organic matter-rich waters (which can promote blooms of heterotrophic organisms leading to oxygen depletion in rivers) sewage treatment deals with a combination of organic particles along with diverse filters and a treatment of the resulting sludge from anaerobic microorganisms (Margot et al 2013).

Similarly, landfills have been widely used as a cheap solution of storing waste, despite the environmental consequences involving pollution on a local scale associated to leaching as well as contributing to global warming due to methane emissions. Some problems associated with the capacity for treatment are related to the presence of heavy metals, organic pollutants (particularly aromatic hydrocarbons) and other problematic components. In order to address these environmental problems, strains of microorganisms could be used to target these molecules.

This approach is known as bioremediation, and has been used with different degrees of success (Cases and de Lorenzo 2005, de Lorenzo 2008). In recent years it has been suggested that the use of genomic search, along with systems and synthetic biology approximations should be considered as a really effective approach to this problem (Schmidt and de Lorenzo 2012, de Lorenzo et al 2016).

If the same basic design used above, namely engineering an existing species, the basic scheme would be summarised in figure 10a-c. To make an abstract and general formulation of the problem, we define a Terraformation motif that incorporates some kind of “container” (grey box) indicating the presence of a boundary condition. This represents for example the sewage system of a urban center or the spatial domain defining the limits of a landfill. In this way, we can define inputs and outputs associated with the inflow of water, organic mater or chemicals on one hand and the outflow carrying other classes of molecules as well as microorganisms on the other.

**FIG. 10.**
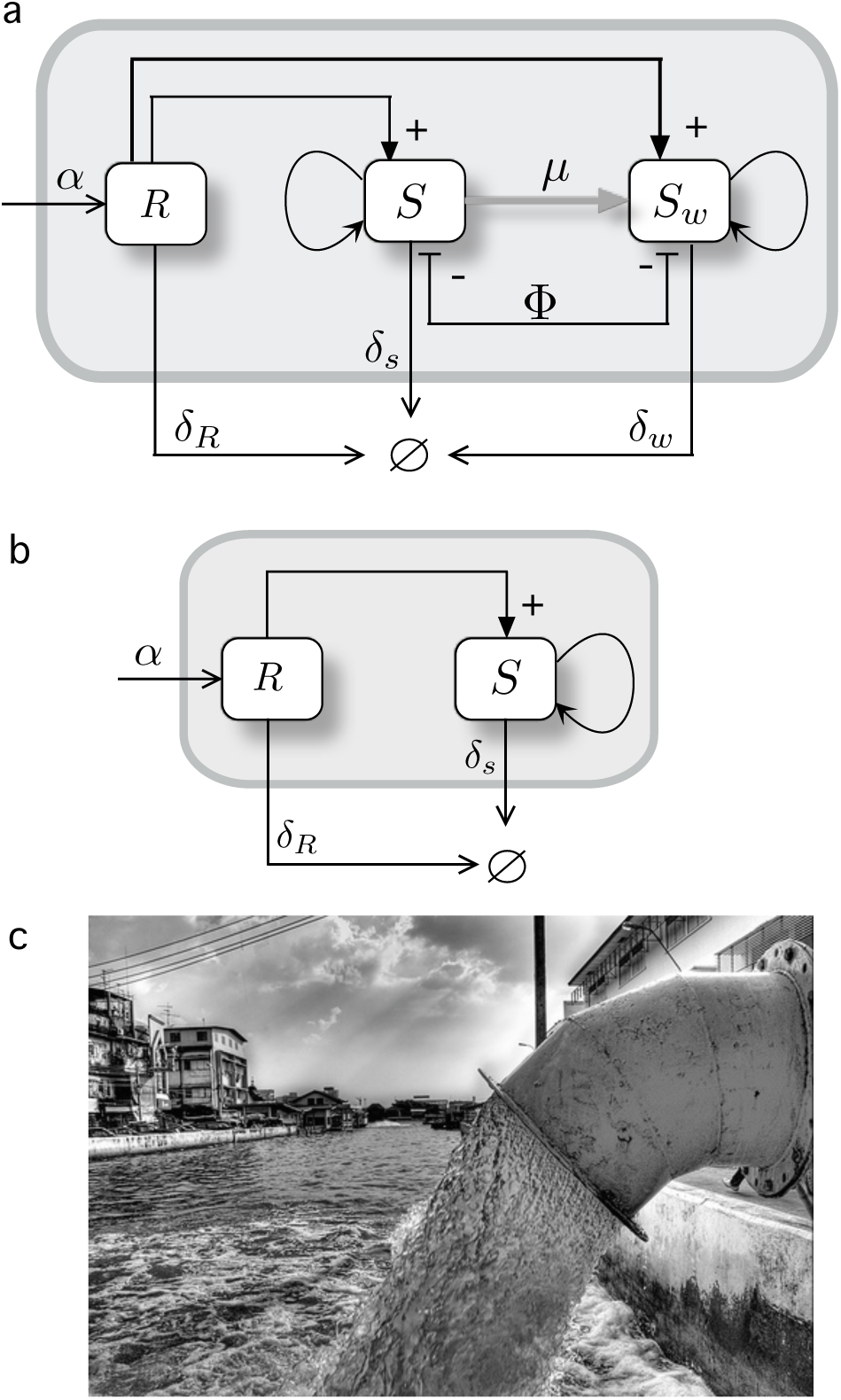
Terraformation motifs for the sewage and landfills. In both (a,b) there exists a a physical container and thus a physical boundary allowing to define input and output flows. In (a) the graph shows the resource-consumer structure of this motif, where both strains are supported by the same resource, while they compete and are connected through gene loss. A simpler alternative (b) does not require engineering of extant species since it is a completely artifactual ecosystem and its preservation is not required. A typical scenario would be sewage-related infrastructures (c) where a rich microbial community is known to exist.

Here too it might be less relevant to preserve the existing species of microbes, given the less relevant motivation of preserving wild type strains, thus making unnecessary to engineering from *S_w_* (figure 10b). Of course the real situation is much complex in terms of species and chemical diversity, and the single box indicating the resource *R* encapsulates a whole universe of chemical reactions. But we can also consider *R* a very specific target for our synthetic strain.

The mathematical model for this motif is given by the next set of equations:

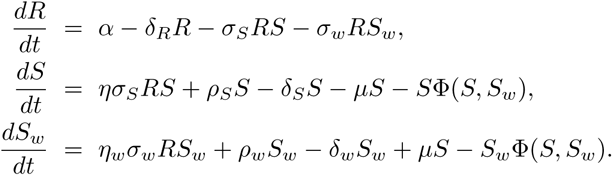

Here, as in the previous model, *δ_R_* denotes the spontaneous degradation rate of the resource. The degradation of the resource by the synthetic and the wildtype strain is parametrized with *σ_S_* and *σ_w_*, respectively. Some fraction of the degraded resource can be invested for growth and reproduction for both the synthetic and the wild-type strains. The constants *η* and *η_w_*, parametrize this process. Assuming constant population the strains, *S* + *S_w_* = 1 and *Ṡ* + *Ṡ_w_* = 0, the competition function is given by Φ(*S, S_w_*) = (*η_w_σ_w_* + *ρ_w_* − *δ_w_*) + *S*(*ησ* − *η_w_* + *ρ͂* − *δ͂*). Using the previous conditions, the three-variable system is reduced to a two-dimensional dynamical system describing the dynamics of the resource (*R*) and the synthetic strain (*S*):

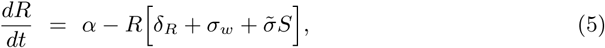

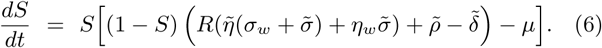

Notice that here, for simplicity, we set *δ͂* = *δ_S_* − *δ_w_*,*ρ͂* = *ρ* − *ρ_w_*,*σ͂* = *σ_S_* − *σ_w_*, and *ῆ* = *η* − *η_w_*.

The fixed points for Eqs. (5)–(6) are given by:

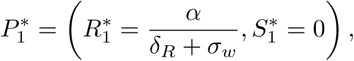

and the pair of fixed points 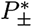 (see Section 3 in the Supplementary Information for their values). The Jacobian matrix the system above reads:

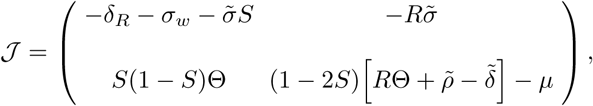

where Θ = [*ῆ*(*σ͂* + *σ_w_*) + *η_w_σ͂*].

The eigenvalues of the first fixed point, obtained from det 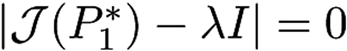, are given by:

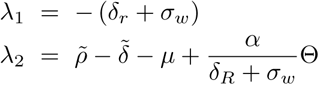

Notice that λ_1_ is always negative, and thus the stability of 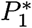 entirely depends on λ_2_. From λ_2_ we can define a critical *μ* value, given by:

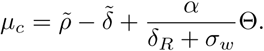

It is easy to see that when *μ* > *μ_c_*, λ_2_ < 0 and thus *P*_1_ is stable. Under this stability condition the synthetic strain will become extinct.

In order to focus on the most interesting parameters from the engineering point of view (i.e., *σ͂* and *ῆ*) we will hereafter take into account that both *S* and *S_w_* strains reproduce at the same rates in the absence of *R* (*ρ͂* = 0), also assuming that both strains have the same death rates (*δ_w_* = *δ_S_* i.e.,*δ͂* = 0). Under these assumption, the equations now read:

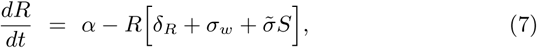

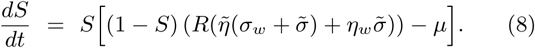

For the system above, the fixed point 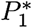 remains the same, but now there exists a single fixed point in the interior of the phase plane. The fixed point is now given by 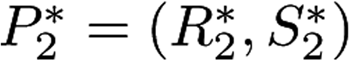 with:

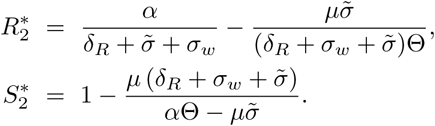

The eigenvalues of the first fixed point, obtained from det 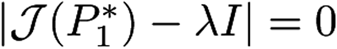, fixing *ρ͂* = *δ͂* = 0, are given by:

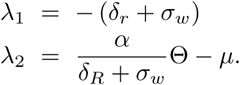

As mentioned above, the stability of this fixed point will depend on λ_2_, and now the critical *μ* value involving a change in the stability of 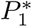 is given by

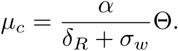

Also, notice that all the rest of the model parameters apart from *μ* are in λ_2_. Hence, the bifurcation can be also achieved tuning these parameters. For the case of *μ*, it can be shown that

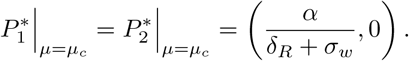

The impact of parameters *σ͂*, *ῆ* and *μ* on the equilibrium concentrations of the resource and the synthetic strain is displayed in figure 11a. We note that the resource in the parameter space (*σ͂, ῆ, μ*) is always present.

**FIG. 11.**
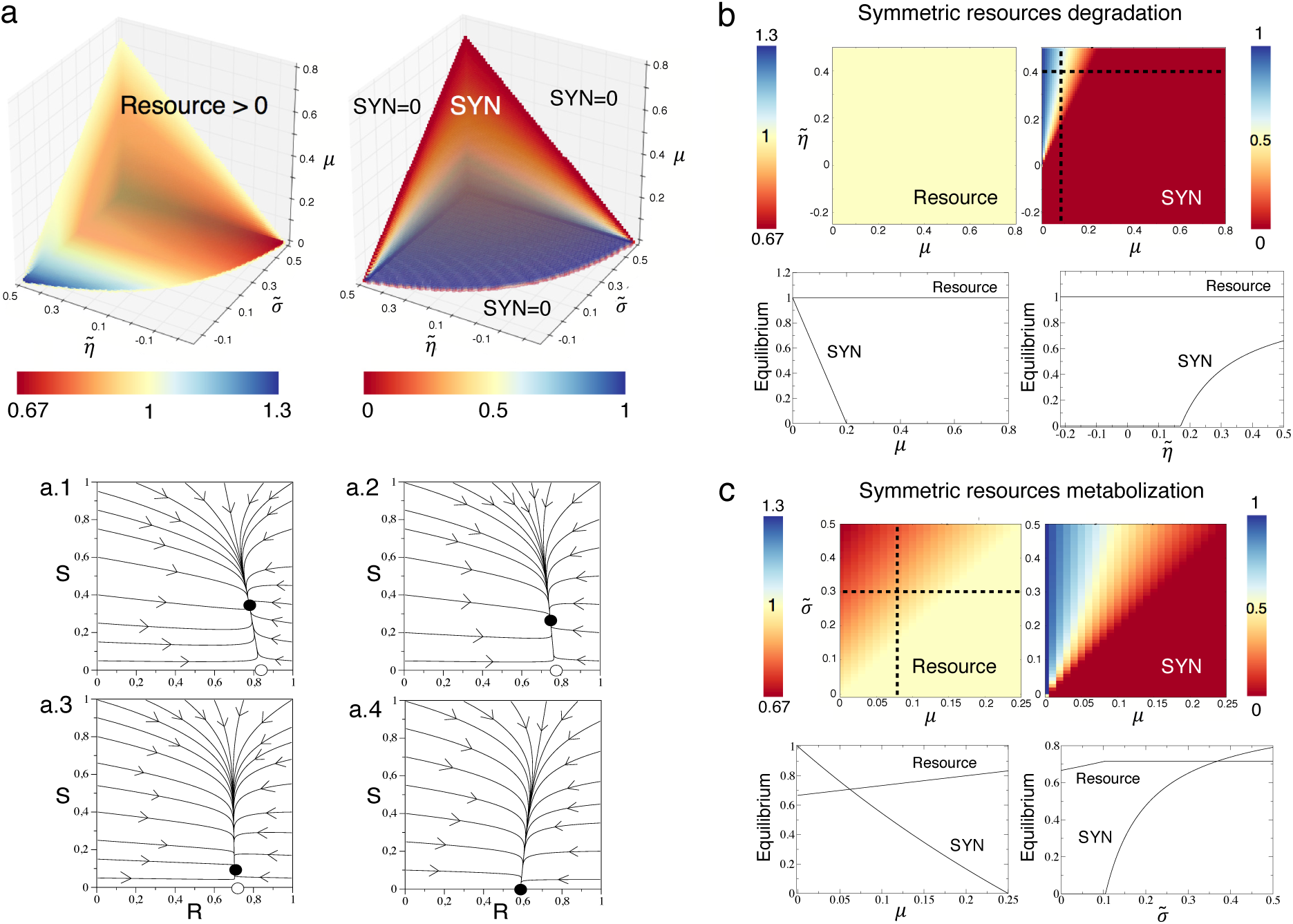
Representative dynamics of the *sewage and landfill motifs*. (a) Equilibrium population values (represented with a color gradient) in the parameter spaces (*σ͂, ῆ, μ*) obtained numerically from Eqs. (7)–(8). We notice that we plot the volume where the synthetic (SYN) strain survives and the resource equilibria for this volume (the resource is always present in the whole space). Below we display several phase portraits with: (a.1) *σ͂* = 0.25, (a.2) *σ͂* = 0.15, (a.3) *σ͂* = 0.03, and (a.4) *σ͂* = −0.07. Here, black and white circles mean stable and unstable fixed points, respectively. The arrows indicate the direction of the flows. (b) Scenario with symmetric resource degradation for both strains, setting *σ_w_* = *σ_S_* ≠ 0 i.e., *σ͂* = 0. We display the equilibria for the resource and the synthetic strain in the parameter space (*μ, ῆ*). Below we display the bifurcation diagrams plotting the population equilibria for both synthetic and resource variables along the parameter values (*μ, ῆ*) indicated with the dashed lines in the panels above. (c) Scenario with symmetric resources metabolization *η_w_* = *η* ≠ 0, with *ῆ* = 0. Here we also plot the population equilibria along the black dashed lines from the panels above in the form of bifurcation diagrams using *μ* and *σ͂* as control parameters. In all of the analysis we set *α* = 1 and *δ_R_* = 0.05.

However, the boundaries causing extinction for the synthetic strain are clearly seen (figure 11a (right)). These transitions are given by the bifurcation previously discussed. Similarly to the previous models, the increase of *μ* involves the extinction of S, being all the population formed by the wild-type strain.

As expected, the increase of both *σ_w_* and *η_w_* also causes the extinction of the synthetic strain. This actually means that if the wild-type has a fitness advantage in terms of resource degradation or metabolization, this population will out-compete the synthetic strain. Notice that this effect takes place when both *σ͂* and *ῆ* decrease (i.e., *σ_w_* and *η_w_* increase, respectively), and the wild-type strain is able to degrade the resource faster or to better metabolize this resource. Under these conditions, and due to the competitive exclusion principle, the synthetic strain will be outcompeted by the wild-type one. This process is accelerated at increasing parameter *μ*. The dynamics tied to different and decreasing values of *σ͂* are represented in the phase portraits of figure 11(a.1-a.4).

Two specific cases can be considered here from the model given by Eqs. (7)–(8) involving two different ecological scenarios that could be achieved by means of different engineering strategies. First, we will consider that both strains degrade the resource at the same rate (symmetric degradation) while their reproduction due to the consumption of such resource can be different. In the second scenario, we will consider that both synthetic and wild-type strains degrade the resource at different rates but the metabolic efficiency of the resource is equal for both strains. Consider first the case were both the wildtype and the synthetic strains degrade the resource at the same rates *σ͂* = 0, with *σ_w_*, ≠0 and *σ_S_* ≠ 0. However, their ability to metabolize the resource and use it for reproduction can be asymmetric (i.e., *ῆ* ≠ 0). The fixed points under these assumptions are given by 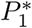 (which remains the same as described above), and:

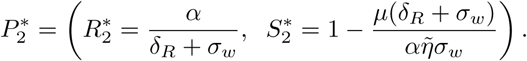

Notice that the bifurcation will take place when 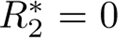 and when λ_2_ = 0. In particular, it can be shown that the synthetic strain will survive when *μ* < *μ_c_*, with:

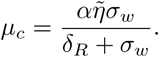

Notice that the diagrams in figure 11(b) indicate such a smooth transition involving the extinction of the synthetic strain. Together with *μ*, the other parameters involved in the transcritical bifurcation are *α*, *ῆ*, *σ_w_*, and *δ_R_*. For instance, the bifurcation diagram tuning *ῆ* in figure 11(b) displays the transition at the value *ῆ* = *ῆ_c_*, with:

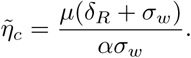

As previously mentioned, the resource is always present, and its population is constant for this case since its equilibrium values does not depend on *μ* and *ῆ*. Notice that the coordinate 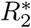 is the same as 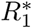, which depend on parameters *α*, *δ_R_*, and *σ_w_*. This is the reason why the equilibrium of the resource does not change at the bifurcation. It is important to highlight that the concentration of the resources will decrease at increasing *σ_w_*. Hence, under symmetric degradation of the resources, the degradation efficiency of the wild-type will determine the equilibrium concentration of the resource. The corresponding bifurcation diagram of figure 11(b) show the transitions of the synthetic strain at *μ* = *μ_c_* (left) and *ῆ* = *ῆ_c_* (right).

Finally, we shall assume that both strains can degrade the resource differently (*σ͂* ≠ 0), but their efficiency to metabolise it (and thus reproduce) is the same, and thus *ῆ* = 0, with *ῆ* ≠ 0 and *η_w_* ≠ 0. Under this scenario, the fixed points are again 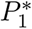, and 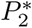 being 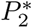 now given by:

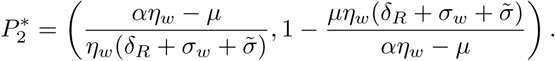

Here, again, the bifurcation values are obtained form λ_2_ computed before. Here, the synthetic strain will also survive when *μ* < *μ_c_*, now with:

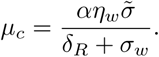

As shown in figure 11(c) the transition takes place at *σ͂* = *σ͂_c_* with:

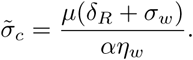

Figure 11(c) shows that the equilibrium concentration of the resource depends on parameters *μ* and *σ͂*, although the concentration of the resource for the parameters analysed remains large. Increasing *σ_w_*, or *μ* (decreasing *σ͂*) involve the extinction of the synthetic strain.

### VIII. DISCUSSION

As we rapidly move to an uncertain future, both ecosystems and societies face the threat of catastrophic responses to a diverse number of external and internal drivers. Several sources of instability are involved in this process, all of which have a direct or indirect anthropogenic origin. The rise of carbon dioxide levels with the inevitable warming of the planet, the always increasing production of waste and a demographic pressure and over-exploitation of constantly shrinking habitats are real challenges that need to be addressed before the inevitable occurs.

Along with other strategies involving the protection of biodiversity hot spots, some geoengineering approaches, sustainable growth and a rational management of resources and non-recyclable waste, particularly when dealing with some key chemicals. But the risk tied to tipping points pose a serious limit to the success of all these strategies. Once unleashed, catastrophic shifts are likely to get amplified by the interconnected web linking ecosystems and essential resources needed to sustain social and economic organization. Moreover, the damage caused by increasing climate variability and drought can trigger social unrest long before any shift occurs (Kelly et al 2015).

To counterbalance the runaway effects derived from the nonlinearities causing shifts we might need to engineer ecological systems. In this context, the proposed framework aims to extend the standard approach of bioremediation (de Lorenzo et al 2016) to larger scales and under a new set of ecological-grounded rules. The models presented here are a first step in defining a population dynamics theory of synthetic ecosystems, that incorporate some classical modelling approaches with a constrain imposed by the nature of the synthetic component (which can revert to a wild type strain).

Humans have been highly successful as engineers, particularly as large-scale ecosystem engineers (Jones et al 1994, 1997) since our activities have deeply modified the energy and matter flows through ecosystems. To a large extent, our common approach here is to design a synthetic microorganism that can modify the ecological interactions in such a way that we engineer an ecological engineer. In fact, it has been already suggested that a useful approach to restoration ecology should consider the major role played by ecosystem engineers and the existence of multiple alternative states (Seastedt 2008, Suding 2004).

All our motifs share a common design principle: the synthetic strain has been derived from a natural one already present within the target community. The aim of this choice is to prevent the failure of the synthetic strain from establishing. Since the engineered cell carries an extra genetic construct, this engineered component will be lost (and reversion will occur) unless the gain in function overcomes the extra metabolic burden. In this paper we have determined the inequalities defining parameter domains where the engineered strains (and their ecological functions) would be stable. The parameter relations derived in the previous sections should help guiding experimental implementations of our Terraformation scenarios.

Because the inevitable simplification imposed by our low-dimensional models, it can be argued that many potential biases might arise from diversity-related factors. Community dynamics might limit or even prevent the spread of the engineered strain, but we also need to consider how the changes derived from the engineering propagate through the system. However, indirect evidence from manipulation experiments inoculation of microorganisms can successfully change the organisation and functionality of a given ecosystem in predictable ways, particularly in relation with soil crust ecosystems (see for example Maestre et al 2006, Bowker 2007, Wu et al 2013). Future work should validate the predictions made here and further explore the limitations and potential extensions of our formalism.

## Acknowledgments

The authors thank the members of the Complex Systems Lab for useful discussions as well as Fernando Maestre and his team for very useful discussions. This study was supported by an European Research Council Advanced Grant (SYNCOM), the Botin Foundation, by Banco Santander through its Santander Universities Global Division and by the Santa Fe Institute. This work has also counted with the support of Secretaria d’Universitats i Recerca del Departament d’Economia i Coneixement de la Generalitat de Catalunya.

